# Scaling of Neuronal Growth and Excitability Through Separable mTORC1 and mTORC2 Pathways

**DOI:** 10.64898/2026.06.22.733785

**Authors:** Asan F Abdulkareem, Noah H Elste, Rain Younger, Mackenzi L Prina, Evan Dennis, Jessica Urbanczyk, Mohsen Bayat, Hassaan A Malik, Shahriar A Tafti, Hunter S Holoubek, Dylan Marchand, Karen E Beenken, Dennis W Province, Kristopher T Kahle, Matthew C Weston, Jeremy M Barry, Bryan W Luikart

**Affiliations:** Department of Neurobiology, Heersink School of Medicine, University of Alabama at Birmingham, Birmingham, AL; Department of Molecular and Systems Biology, Geisel School of Medicine, Dartmouth College, Hanover, NH; Department of Neurological Sciences, Larner College of Medicine, University of Vermont, Burlington VT; Fralin Biomedical Research Institute, Virginia Tech, Roanoke, VA; IDEA National Resource for Quantitative Proteomics, University of Arkansas for Medical Sciences, Little Rock, AR; Department of Neurosurgery, Massachusetts General Hospital & Harvard Medical School, Boston, MA

## Abstract

As neurons grow, they must regulate intrinsic excitability to maintain an appropriate level of spiking based on synaptic inputs. Using gene knockouts, phosphoproteomics, and electrophysiology we show PTEN regulates neuronal growth and intrinsic excitability through separable downstream mechanisms. *Pten* loss induces cellular hypertrophy, increased excitatory synaptic input, reduced fast afterhyperpolarization, and burst firing. Deleting the mTORC1 scaffold, *Raptor,* rescues overgrowth and synaptic input but fails to normalize firing, while deleting *Akt* or the mTORC2 scaffold, *Rictor,* restores firing without rescuing growth. This dissociation identifies an AKT-mTORC2 mechanism that regulates voltage-gated calcium and BK potassium channels to set spike repolarization and burst firing. *In vivo*, *Pten* knockout produces altered network synchrony, lethal seizures, and impaired object and location behavior; *Raptor* co-deletion display non-lethal hyperexcitability with improved object-location coupling. The biological and pathophysiological significance of these mechanisms is demonstrated by overlap of the PTEN-regulated phosphoproteome with ASD and epilepsy.

## Introduction

Neurons must coordinate their growth with their excitability to generate appropriate responses to synaptic input (Henneman et al., 1965; O’Leary et al., 2014, 2015). As a developing neuron elaborates its axonal and dendritic processes, the surface area available for synaptic input grows, the membrane capacitance increases, and the input resistance falls. Larger neurons therefore contain more ion channels and produce larger aggregate currents, yet a given synaptic current generates a smaller voltage response. To maintain functional output through this developmental enlargement, neurons must coordinately scale their voltage-gated ion channel complement and their synaptic input strength to match their changing morphology (Marder and Goaillard, 2006; Turrigiano and Nelson, 2004). Failure to coordinate these processes produces neurons that are either electrically silent (large cells with insufficient channel density or input) or pathologically excitable (cells with channel composition mismatched to their geometry). How neurons solve this scaling problem during development, and what molecular machinery couples growth to intrinsic excitability, remains incompletely understood (Blankenship and Feller, 2010; Yuste et al., 2024).

The PI3K/PTEN signaling axis is well-suited to coordinate these processes. Receptor tyrosine kinases activate PI3K to generate PIP3, and PTEN reverses this reaction by dephosphorylating PIP3 to PIP2. PIP3 recruits AKT to the membrane and engages two distinct mTOR complexes. mTORC1, defined by the scaffold Raptor and inhibited by rapamycin, drives cap-dependent protein translation and cellular growth. Canonically, mTORC2, defined by Rictor, phosphorylates AKT to augment mTORC1 activity and independently regulates cytoskeletal remodeling. In neurons, loss of PTEN engages both complexes and produces a stereotyped constellation of phenotypes including cellular hypertrophy, dendritic and synaptic overgrowth, reduced afterhyperpolarization, and increased burst firing (Kwon et al., 2006; Luikart et al., 2011; Santos et al., 2017; Williams et al., 2015). PTEN mutations cause PTEN Hamartoma Tumor Syndrome (PHTS), a pleiotropic disorder characterized by macrocephaly, benign tumors, cancer, autism spectrum disorder (ASD), epilepsy, and hydrocephalus (Dhawan et al., 2024; Macken et al., 2019; Yehia and Eng, 1993). Disorder-focused whole-exome sequencing demonstrates that *PTEN* is among the most commonly mutated genes in patients with ASD (Fu et al., 2022), the most commonly mutated gene in patients with ventriculomegaly or hydrocephalus (DeSpenza et al., 2025), and is associated with epilepsy (Chen et al., 2024).

Whether the morphological and excitability consequences of PTEN loss reflect a single coordinated program or the engagement of independently regulable mechanisms is unknown (Barrows et al., 2017; Skelton et al., 2020). Pharmacological or genetic suppression of mTORC1 with rapamycin or *Raptor* deletion reverses neuronal hypertrophy and synaptic overgrowth (Getz et al., 2016; Narvaiz et al., 2023; Tariq et al., 2022; Zhou et al., 2009), yet seizures and cognitive deficits persist in patients and animal models treated with mTORC1 inhibitors (Getz et al., 2016; Narvaiz et al., 2023; Srivastava et al., 2022; Tariq et al., 2022; Zhou et al., 2009). Recent work has implicated mTORC2 signaling in seizure activity that persists despite mTORC1 inhibition (Chen et al., 2019; Cullen et al., 2024; Okoh et al., 2023), but the contributions of mTORC1 and mTORC2 to neuronal excitability have not been cleanly separated, and the downstream substrate networks they control in neurons remain incompletely defined.

Here, we exploit the experimentally accessible dentate granule neuron using retroviral and transgenic conditional knockout strategies to dissect PTEN signaling. We show that mTORC1 and AKT-mTORC2 control molecularly and functionally distinct cellular programs. mTORC1 drives neuronal growth and excitatory synaptic input, while AKT-mTORC2 regulates the phosphorylation and function of voltage-gated calcium and BK potassium channels that determine action potential repolarization and burst firing. Genetic restoration of normal cellular size and synaptic input through *Raptor* co-deletion does not rescue the AKT-mTORC2-dependent ion channel program. Critically, *Raptor* deletion in *Pten* knockout neurons produces cells with wild-type capacitance and excitatory synaptic drive, isolating AKT-mTORC2-dependent ion channel dysregulation from cell size or input strength. *In vivo*, this dissociation manifests as altered network synchrony, electrographic seizures, and improved object/place coupling behavior, even when the *Pten*-mediated overgrowth is suppressed. These findings reframe neuronal mTORC1 and mTORC2 signaling: mTORC1 scales neuronal growth and synaptic input, while AKT-mTORC2 tunes intrinsic excitability, demonstrating that the coordinated scaling of growth and excitability is regulated through molecularly separable programs. The biological importance of the coordinated regulation of neuronal growth and excitability is substantiated by the overlap of phosphoproteomic profiling of signaling dysregulation downstream of PTEN and high-confidence ASD and epilepsy risk genes.

## Results

### Phosphoproteomics Show mTORC1 Dependent Growth Regulation and mTORC2 Dependent Ion Channel Regulation

We have previously demonstrated that *Pten* KO results in neuronal hypertrophy and hyperexcitability (Luikart et al., 2011; Weston et al., 2014; Williams et al., 2015). We found that ablation of mTORC1 via *Raptor* KO completely rescued all morphological overgrowth including increased spine density and excitatory synapse number while mTORC2 inhibition via *Rictor* KO partially rescued morphological overgrowth and synapse formation caused by *Pten* KO (Cullen et al., 2023; Tariq et al., 2022). However, neither the *Pten;Raptor* DKO, nor the *Pten;Rictor* dKO was able to stop seizures in these mice (Cullen et al., 2024). To determine the mechanism leading to hyperexcitability in these dKOs, we performed a phosphoproteomic analysis on the dentate gyrus from *Pten^flx/flx^* x POMC-Cre+ (*Pten* KO) and *Pten^flx/flx^;Raptor^flx/flx^* x POMC-Cre+ (*Pten;Raptor* dKO) mice. POMC-Cre is expressed in most of the dentate granule neurons with higher percentages in dorsal than ventral dentate gyrus (Gonzalez et al., 2023).

Using tandem mass tag labeling (TMT-11), we performed multiplex phosphoproteomic analysis of phosphopeptides regulated in *Pten^flx/flx^* x POMC-Cre+ (n=4) and *Pten^flx/flx^;Raptor^flx/flx^*x POMC-Cre+ (n=4) relative to *POMC-Cre*- (wild-type, n=3) controls. Comparing *Pten* KO versus wild-type with log fold change > ±1.1 and p < 0.05, we identified 1,737 differentially regulated phosphopeptides corresponding to 901 proteins (Figure 1A). For *Pten;Raptor* dKO mice, we identified 469 phosphopeptides corresponding to 319 proteins (Figure 1B). Of these 319 *Pten;Raptor* dKO-regulated proteins, 301 overlapped with the set of 901 proteins regulated in *Pten* KO (Figure 1C). This is consistent with the interpretation that *Pten* KO is the major driver of physiological change and that co-deletion of *Raptor* rescues the portion of those changes related to translation and overgrowth.

**Figure 1.**
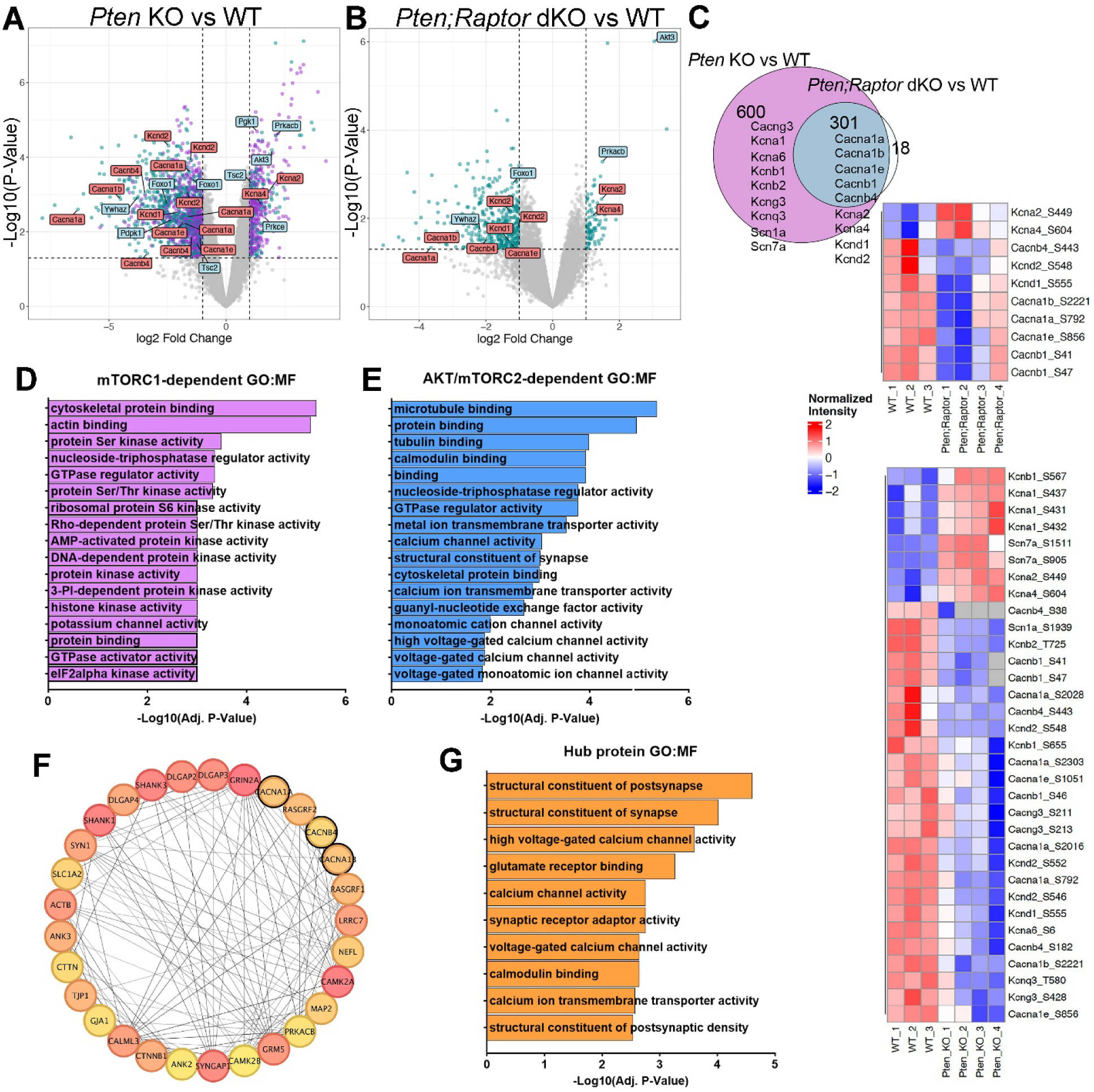
Phosphoproteomic Analyses Reveal Network of mTORC1-dependent Neuronal Growth and Akt-mTORC2-dependent Excitability. (A) Volcano plot showing peptides with differential phosphorylation between *Pten* KO and wild-type samples. Peptides significantly regulated only in *Pten* KO are shown in purple; peptides significantly regulated in both *Pten* KO and *Pten;Raptor* dKO are shown in green. Peptides corresponding to “ground truth” proteins upstream of mTORC1 are labeled in blue, and differentially regulated voltage-gated ion channels are labeled in red. (B) Volcano plot of differentially phosphorylated peptides in *Pten;Raptor* dKO versus wild-type. “Ground truth” upstream mTORC1 proteins are labeled in blue and voltage-gated ion channels in red. (C) Proportional Venn diagram showing overlap of differentially phosphorylated proteins in *Pten* KO versus wild-type (901 peptides, purple) and *Pten;Raptor* dKO versus wild-type (319 peptides, green). Of the 319 peptides regulated in *Pten;Raptor* dKO, 301 overlap with the *Pten* KO dataset. Voltage-gated ion channels identified in *Pten* KO alone or in both genotypes are listed. Heatmaps display normalized intensities of voltage-gated ion channel peptides regulated in *Pten;Raptor* dKO (top) and *Pten* KO (bottom). (D) Bar plot of the top Gene Ontology Molecular Function (GO:MF) terms enriched among mTORC1-dependent phosphoproteins regulated only in *Pten* KO, ranked by −log_10 adjusted *P* value. (E) Bar plot of top GO:MF terms enriched among AKT/mTORC2-dependent proteins regulated in both *Pten* KO and *Pten;Raptor* dKO, ranked by −log_10 adjusted *P* value. (F) Protein–protein interaction (PPI) network of the top 29 of 30 hub proteins with highest connectivity among the 301 proteins regulated in both *Pten* KO and *Pten;Raptor* dKO. Darker node fill indicates higher centrality; voltage-gated ion channels are outlined in black. (G) Bar plot of enriched GO:MF terms among hub proteins shown in F.

Among the regulated phosphoproteins that remained in the *Pten;Raptor* dKO were 15 voltage-gated Ca²⁺ and K⁺ channel proteins. Voltage-gated Ca²⁺ channels were 12x enriched in the mTORC1-independent set (Fisher OR = 12.12, p = 0.007). Altered phosphorylation of voltage-gated Ca²⁺ channel subunits in *Pten;Raptor* dKO mice included CACNA1A (P/Q-type), CACNA1B (N-type), CACNA1E (R-type), and CACNB1 and CACNB4 (L-type) (Figure 1A-C). We also observed altered phosphorylation of K_V_1 and K_V_4 voltage-gated K^+^ channels. Further, every ribosomal protein and translation initiation factor was exclusive to the 600 proteins regulated only in the *Pten* KO (RPLP0, RPLP1, RPLP2, RPS6KA4, RPS6KC1, RRBP1, EIF4G1, EIF5B).

We next performed gene ontology analysis of molecular function to assess the biological activities enriched within each phosphoproteomic program, using the 6,446-protein union dentate gyrus proteome as the enrichment background. Among the 600 proteins regulated exclusively in *Pten* KO mice, the top enriched GO Molecular Function terms included protein serine/threonine kinase activity (GO:0004674, p = 4.75 × 10⁻⁴), ribosomal protein S6 kinase activity (GO:0004711), and eIF2α kinase activity (GO:0004694), consistent with broad activation of the mTORC1-driven translational program (Figure 1D). In contrast, the 301 proteins regulated in both *Pten* KO and *Pten;Raptor* dKO were enriched for calmodulin binding (GO:0005516), calcium channel activity (GO:0005262), calcium ion transmembrane transporter activity (GO:0015085), and voltage-gated calcium channel activity (GO:0005245), including high voltage-gated calcium channel activity (GO:0008331) (Figure 1E). None of these calcium channel terms reached significance in the mTORC1-dependent set. To further define relationships among the 301 AKT/mTORC2-regulated proteins, we conducted protein-protein interaction analysis using the STRING database and identified highly connected hub proteins. Twenty-nine of the top 30 hub proteins formed a dense interaction network whose most enriched GO Molecular Function terms were structural constituent of postsynapse, structural constituent of postsynaptic density, high voltage-gated calcium channel activity (GO:0008331), glutamate receptor binding, calcium channel activity (GO:0005262), voltage-gated calcium channel activity (GO:0005245), and calmodulin binding (GO:0005516), and included the voltage-gated calcium channel subunits CACNA1A, CACNA1B, and CACNB4 (Figure 1F,G).

### mTORC1-Independent Regulation of Neuronal Burst Firing by PTEN

To determine the physiological impact on neuronal excitability after *Pten* loss, we used a combination of mouse floxed alleles and dentate gyrus (DG) *in vivo* retroviral injections at P7 to express Cre recombinase and fluorophores to label, birthdate, and knockout floxed alleles in developing neurons (Figure 2A)(Cullen et al., 2023; Tariq et al., 2022; Williams et al., 2015).

**Figure 2.**
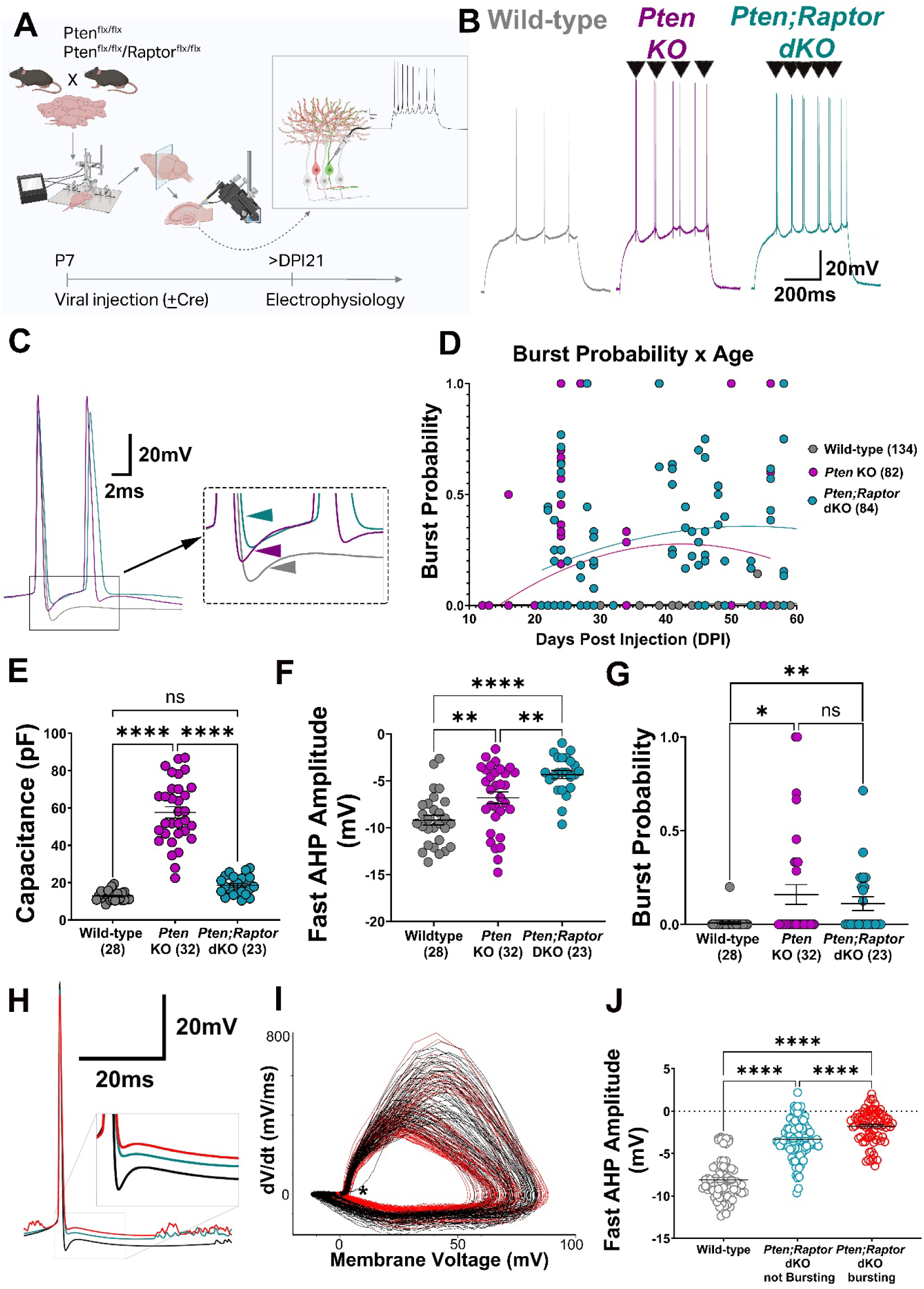
*Pten* and *Pten*;*Raptor* KO Neurons Display Burst-Firing with Decreased fAHP. (A) Stereotaxic injection of *Pten^flx/flx^*or *Pten^flx/flx^;Raptor^flx/flx^* mice with Cre expressing retroviruses at P7 was used to label, birthdate, and knockout the respective floxed genes in developing dentate gyrus granule neurons. (Created in Biorender (Licensed): https://biorender.com/tfsc7bw). (B) Sample whole-cell current-clamp traces of wild-type, *Pten* KO, and *Pten;Raptor* dKO neurons responding to a threshold 500ms depolarization. Bursts of 2 action potentials firing within 20ms are indicated by black arrowheads. (C) Close-up of the first action potential in wild-type (gray), *Pten* KO (magenta), and *Pten;Raptor* dKO (cyan) show the two action potential burst in the *Pten* KO and *Pten; Raptor* dKO. The inset displays the decreased fast afterhyperpolarization potential (fAHP) amplitude in the *Pten* KO and *Pten;Rapto*r dKO (color arrows). (D) A plot of action potential burst probability versus neuron age, assayed as the number of days post-retrovirus injection (DPI) indicates that mature *Pten* KO and *Pten;Raptor* dKO neurons display increased burst-firing compared to wild-type. (E) Quantitative data (mean±SEM) shows that the *Pten* KO increases neuronal capacitance and *Pten*;*Raptor* dKO rescues capacitance. (each point is a neuron, n, is indicated in parentheses; *p<0.05, **p<0.01, ***p<0.001 versus wild-type; one-way ANOVA with Bonferroni post-hoc) (F) Quantitative data (mean±SEM) shows that the *Pten* KO and *Pten;Raptor* dKO decrease the amplitude of the fAHP. (G). Plot showing that *Pten* KO and *Pten;Raptor* dKO increase action potential burst probability. (*p<0.05, **p<0.01, Fisher’s exact test). (H) Average traces generated by capturing the first action potential from the first threshold current step (rheobase) that was not followed by a second action potential for 20ms from wild-type (black), *Pten;Raptor* dKO neurons not displaying burst-firing (cyan), and bursting *Pten;Raptor* dKO (red). (I) Overlaid phase plots for all wild-type (black), and *Pten;Raptor* dKO (red) action potentials captured at rheobase depict a consistent alteration in the fAHP (asterisk). (J) *Pten;Raptor* dKO neurons have a decreased AHP amplitude compared to wild-type and the magnitude of that decrease could differentiate between bursting and non-bursting *Pten;Raptor* dKO neurons. Each point represents a single action potential (*p<0.05, **p<0.01, ***p<0.001 versus wild-type; one-way ANOVA model with Bonferroni post-hoc).

This allowed us to examine the physiological parameters of KO neurons using wild-type neurons from the same animals as a control to isolate the impact of gene knockout from any uncontrolled between-mouse variability. We examined electrophysiological parameters in wild-type (fluorophore-expressing), *Pten^flx/flx^*, Cre+ (*Pten* KO) and *Pten^flx/flx^*; *Raptor^flx/flx^*, Cre+ (*Pten;Raptor* dKO) DG granule neurons between 24-60 days post-injection (Figure 2A). We found that both *Pten* KO and *Pten;Raptor* dKO display prominent burst-firing of two or more action potentials within 20ms at rheobase (Figure 2B) with no change in spike amplitude. In addition to increased burst firing, we examined the spike waveform of the first action potential fired at rheobase and found a decrease in the peak amplitude of the fast afterhyperpolarization potential (fAHP) (Figure 2C). By analyzing recordings from younger neurons from 7-25 days post-injection (recorded Williams et al 2025) and the present set we determined the developmental onset of bursting occurs after 21 days post-injection in mature neurons (Figure 2D). The *Pten;Raptor* dKO exhibit cellular capacitance indistinguishable from wild-type, confirming the morphological rescue of neuronal size seen in Tariq et al. 2022 (Figure 2E). However, despite the morphological rescue, both decreased fAHP and high burst probability persisted in the *Pten;Raptor* dKO (Figure 2F,G). Altogether, these data indicate that this form of hyperexcitability is independent of mTORC1.

Because wild-type and *Pten;Raptor* dKO neurons did not differ in membrane capacitance or input resistance, detailed action potential properties were compared between these groups. Spike and afterhyperpolarization (AHP) kinetics were analyzed by isolating all action potentials elicited at rheobase (the first current step at which spiking occurred) and excluding spikes followed by a second action potential within 20 ms, thereby allowing uninterrupted measurement of the AHP (Figure 2H). Phase plots (dV/dt) demonstrated more variability in spike amplitude and rise-time that overlapped between groups with a significant difference in AHP amplitude (Figure 2I). *Pten;Raptor* dKO neurons exhibited reduced fast AHP amplitude and half-width relative to wild-type controls, with a more pronounced decrease observed in bursting *Pten;Raptor* dKO neurons compared to non-bursting *Pten;Raptor* dKO neurons (Figure 2J). Overall, there is an increase in the burst probability and a decrease in fAHP amplitude in both *Pten* KO and *Pten;Raptor* dKO neurons and those cells that fired bursts also had the smallest fAHP amplitudes.

### AKT and mTORC2 are Necessary for Burst Firing of *Pten* KO Neurons

We next asked whether knockout of signaling intermediates in combination with *Pten* could rescue fAHP amplitude and burst probability. We targeted 30-50 day old neurons where bursting was most robust from the time course in Figure 2D and compared to neighboring uninfected wild-type neurons. This independent set of *Pten;Raptor* dKO neurons exhibited increased burst probability accompanied by reduced fAHP amplitude (Figure 3A). Both *Pten;Raptor* dKO and *Pten;Akt1;Akt3* triple knockout (tKO) neurons restored the elevated membrane capacitance observed in *Pten* KO neurons to near wild-type levels, whereas the *Pten;Rictor* dKO did not (Figure 3B). These results align with previous studies demonstrating that loss of either *Raptor* or *Akt1/3* is sufficient to rescue neuronal hypertrophy, whereas Rictor loss is not (Cullen et al., 2024; Cullen et al., 2023; Prina et al., 2026; Tariq et al., 2022). None of the genotypes altered spike amplitude (Figure 3C). In contrast, both the *Pten;Rictor* dKO and the *Pten;Akt1;Akt3* tKO restored the fAHP amplitude and reduced burst probability to levels indistinguishable from wild-type neurons (Figure 3D,E). Together, these data support a model where AKT and mTORC2 signaling decrease fAHP amplitude and promote neuronal burst firing (Figure 3F).

**Figure 3.**
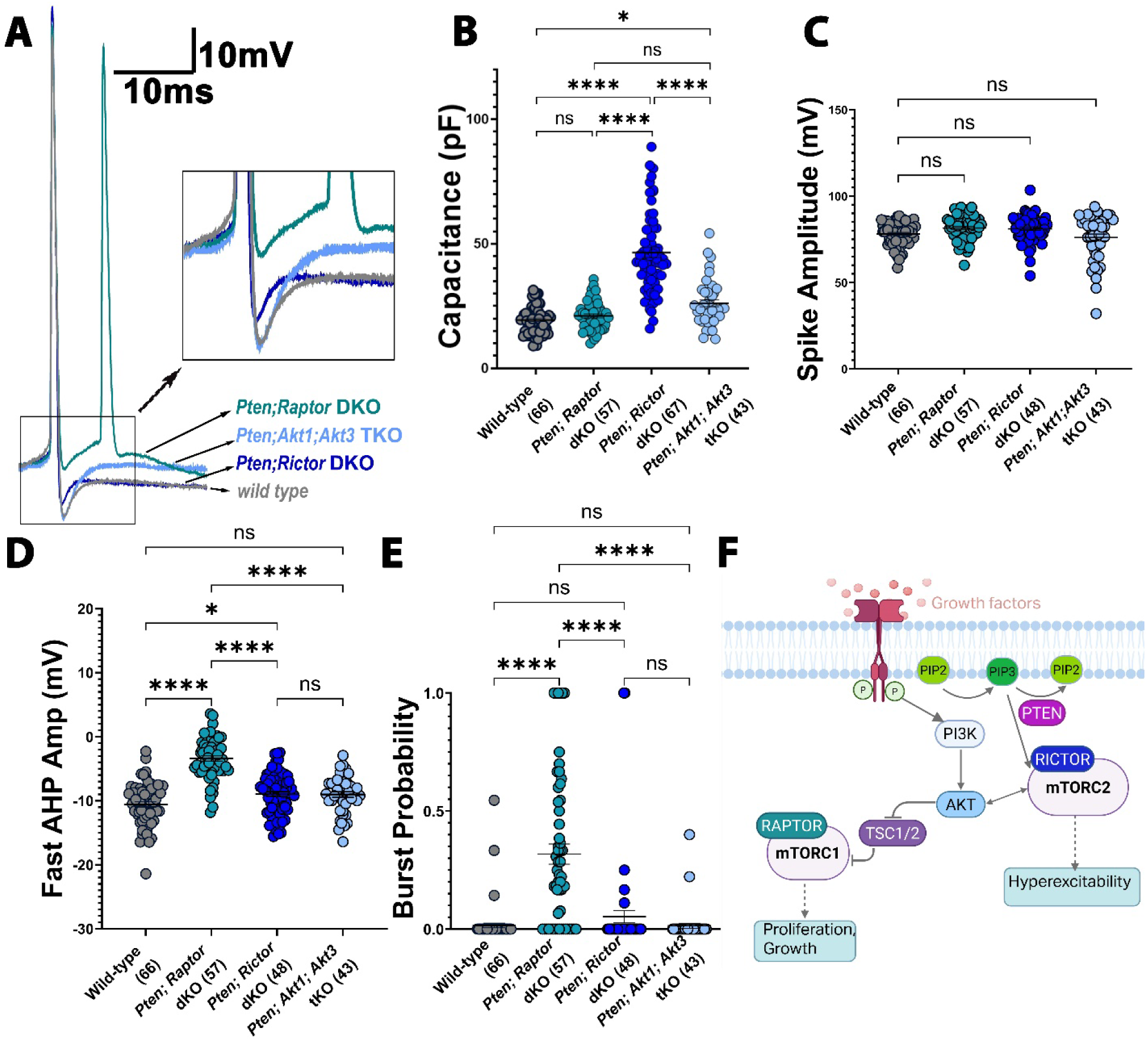
AKT and mTORC2 are Necessary for Burst-Firing. (A) Representative whole-cell current-clamp traces of the first action potential in wild-type, *Pten;Raptor* dKO, *Pten;Rictor* dKO, and *Pten;Akt1;Akt3* tKO neurons in response to a 500 ms depolarizing current injection. *Pten;Rictor* dKO and *Pten;Akt1;Akt3* tKO neurons show increased fast afterhyperpolarization (fAHP) amplitude and reduced burst firing (inset). (B) Quantification indicates that *Pten;Raptor* dKO and *Pten;Akt1;Akt3* tKO neurons display near wild-type cellular capacitance, whereas *Pten;Rictor* dKO neurons exhibit increased capacitance. (C) Average spike amplitude is unchanged across genotypes. (D,E) Fast AHP amplitude (D) and burst probability (E) are restored to wild-type levels in *Pten;Rictor* dKO and *Pten;Akt1;Akt3* tKO neurons. Each dot represents one neuron; (*p<0.05, **p<0.01, ***p<0.001, ****p<0.0001 one-way ANOVA with Tukey’s for D and Fisher’s exact test for E). (F) Schematic of receptor tyrosine kinase signaling in neurons illustrating how the balance between PTEN and PI3K regulates PIP3 production to activate AKT directly and via PDK (not shown). AKT and PIP3 are required for mTORC2 activation, which augments AKT activity in a positive feedback loop. AKT relieves inhibition of mTORC1 through TSC and RHEB (not shown). mTORC1 promotes increased neuronal growth and capacitance, whereas mTORC2 regulates ion channel function and burst-firing hyperexcitability.

### Decreased BK and Voltage-Gated Calcium Currents Underly Bursting in *Pten* and *Pten; Raptor* dKO

Voltage-gated ion channels shape action potentials and the AHP. To define the mechanisms underlying the altered action potential firing, we screened for changes in ionic currents in wild-type versus *Pten;Raptor* dKO neurons. To determine which ionic conductances contributed to the reduced fast AHP amplitude in the *Pten; Raptor* dKO neurons we first examined voltage-gated potassium channel function since previous studies showed a correlation between these channels and bursting (Irie and Trussell, 2017; Niday and Bean, 2021). We observed a reduction in the aggregate voltage-gated K^+^ current in the *Pten; Raptor* dKO (Figure 4A,B,C, Figure S1). To determine if specific channel subtypes contributed to this reduction, we screened for the effectiveness of potassium channel blockers to reduce the voltage-gated potassium currents in wild-type versus *Pten;Raptor* dKO neurons. Paxilline (3µM), an inhibitor of K_Ca_1.1(BK) channels encoded by the *Kcnma1* gene displayed a prominent block in wild-type cells that was decreased in *Pten; Raptor* dKO and in *Pten* KO neurons (Figure 4D,E,F and Figure S1). There was no significant change in the percentage of block of this current by 100nM α-dendrotoxin (K_V_1), 100nM guangxitoxin (K_V_2), 100µM XE991 (K_V_7), or 10µm ML252 (K_V_7.2) (Figure S1). While these experiments do not rule out small changes in currents from other K^+^ channels, they demonstrate a prominent decrease in BK current in *Pten; Raptor* dKO and in *Pten* KO neurons.

**Figure 4.**
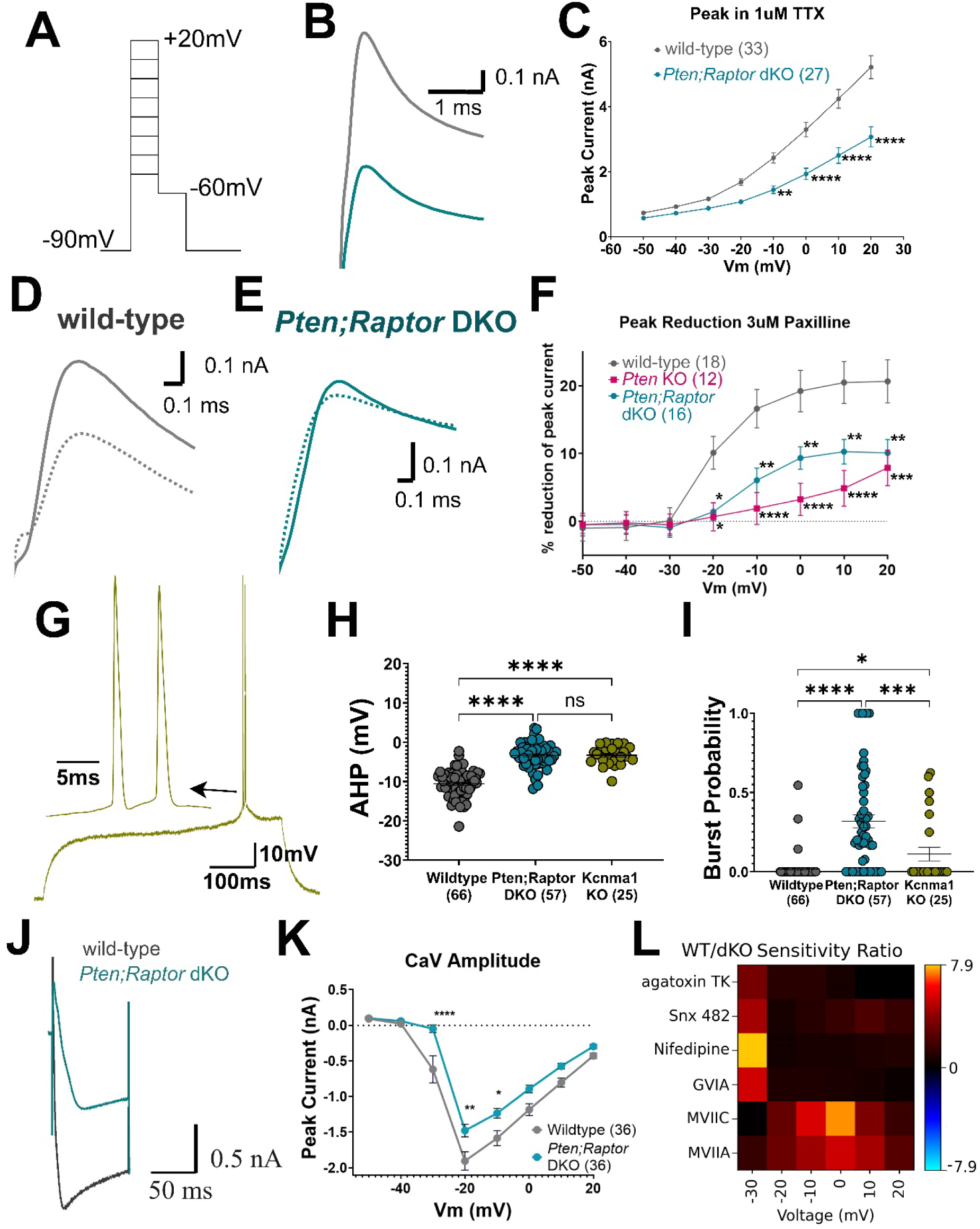
BK and CaV Currents are Reduced in *Pten KO* and *Pten;Raptor dKO*. (A) Voltage-clamp protocol used to examine voltage-gated K^+^ currents. Neurons were held at −90 mV and stepped from −50 to +20 mV for 50 ms. (B) Representative current traces showing peak K^+^ current amplitude at +20 mV in a representative wild-type (gray) and *Pten;Raptor* dKO (cyan) neurons. (C) Quantification of peak K^+^ current amplitude (mean ± SEM) across voltage steps reveals reduced voltage-gated K^+^ currents in *Pten;Raptor* dKO neurons. n values (neurons) are indicated in parentheses; (*p<0.05, **p<0.01, ***p<0.001, ****p<0.0001 versus wild-type using a mixed-effects model with multiple comparisons). (D) Representative recording from a wild-type neuron showing that paxilline application (3 µM, dashed line) reduces peak K^+^ current at +20 mV. (E) Representative recording from a *Pten;Raptor* dKO neuron (cyan) demonstrating minimal response to paxilline application (3 µM, dashed line). (F) Quantification of percent reduction in peak K^+^ current (mean ± SEM) following paxilline application in wild-type (gray), Pten KO (magenta), and *Pten;Raptor* dKO neurons. (*p<0.05, **p<0.01, ***p<0.001, ****p<0.0001, versus wild-type using mixed-effects model with multiple comparisons). (G) Representative recording from a Kcnma1 KO neuron exhibiting burst firing at rheobase. (H) Peak fAHP amplitude is reduced to a similar extent in *Pten;Raptor* dKO and Kcnma1 KO neurons compared to wild-type. n values (neurons) are indicated in parentheses; ****p<0.0001; one-way ANOVA with Tukey’s multiple comparison test. (I) Burst probability for wild-type, *Pten;Raptor* dKO, and Kcnma1 KO neurons. (*p<0.05, **p<0.01, ***p<0.001, ****p<0.0001 using Fisher’s exact test). (J) Representative current traces showing peak Ca^2+^ current amplitude at −20 mV in wild-type (gray) and *Pten;Raptor* dKO (cyan) neurons. (K) Quantification of peak Ca^2+^ current amplitude (mean ± SEM) across voltage steps demonstrates reduced voltage-gated Ca^2+^ currents in *Pten;Raptor* dKO neurons. n values (neurons) are indicated in parentheses; *p<0.05, **p<0.01, ***p<0.001, ****p<0.0001, versus wild-type using mixed-effects model with multiple comparisons) (L) Heatmap illustrating differences in sensitivity of CaV currents to channel blockers (y-axis) across voltage steps (x-axis). Bright orange indicates voltages at which drug application more effectively reduced Ca^2+^ current in wild-type compared to *Pten;Raptor* dKO neurons.

To confirm the role of BK currents in bursting and fAHP, we examined action potentials in *Kcnma1^flx/flx^* mice infected with Cre-expressing retrovirus. *Kcnma1* KO neurons exhibited a decreased fAHP amplitude equivalent to that of *Pten;Raptor* dKO (Figure 4G,H). We also found 6 of 25 *Kcnma1* KO neurons displayed burst-firing. While significant, this frequency was less than that seen in the *Pten;Raptor* dKO (Figure 4G,I). Thus, loss of BK current produced a decreased fAHP amplitude indistinguishable from the *Pten* KO and *Pten;Raptor* dKO but did not fully replicate the burst firing caused by *Pten* KO. This may indicate that BK loss alone is not sufficient to phenocopy the altered excitability of *Pten* KO neurons.

Because BK current depends on both membrane voltage and intracellular calcium concentration and the proteomic data showed a prominent signature of voltage-gated Ca^2+^ channel regulation, we performed a series of experiments examining voltage-gated Ca^2+^ currents (CaV). We found the aggregate (total) CaV current amplitude was decreased in the *Pten;Raptor* dKO (Figure 4J-K). There was a positive shift in the voltage dependence of Ca^2+^ current noted when stepping from -90mV to -30mV producing almost no current in *Pten;Raptor* dKO (46±53pA) compared to wild-type (617±190pA) (Figure 4K). We next examined whether there was differential block of the peak amplitude in response to specific CaV inhibitors. We found evidence for dysregulation of multiple voltage gated Ca^2+^ channels. For channels opening when the neurons were stepped to -30mV wild-type neurons displayed current reduction to 10µM nifedipine (L-type), 1µM ω-conotoxin GVIA(N-type), 0.5µM Snx482 (R-type), and 0.2µM AgatoxinTK (P/Q-type) that was not seen in dKO neurons (Figure 4L, Figure S2).

The decreased response to blockers at -30mV could be due to differential block or it may reflect that there is little to no Ca^2^ current at -30mV for the *Pten;Raptor* dKO neurons. There was differential blockade of current amplitude of *Pten;Raptor* dKO neurons compared to wild-type in response to the -20, -10 and 0mV steps for 1µM ω-conotoxin MVIIC (P/Q N-type) and 0.5µM MVIIA (N-type) (Figure 4L, Figure S2). Overall, and congruent with phosphoproteomics, we detected an altered response for *Pten;Raptor* dKOs compared to wild-type to L, R, P/Q, and N-type channel blockers. We conclude that voltage-gated Ca^2+^ currents and voltage and Ca^2+^-sensitive BK currents are major contributors to altered fAHP amplitude and burst firing in the *Pten* and *Pten;Raptor* KOs.

### *Pten* KO and *Pten;Raptor* dKO Have Hyperactive Dentate Gyri *In Vivo*

We next examined whether the altered action potential firing observed in acute slices from *Pten* KO and *Pten;Raptor* dKO resulted in changes in neuronal circuit function *in vivo*. We injected P7 *Pten^flx/flx^* or *Pten^flx/flx^;Raptor^flx/flx^*mice with retroviruses expressing only GFP (wild-type) or retroviruses expressing Cre (*Pten* KO or *Pten;Raptor* dKO, respectively). Laminar probes in head-fixed mice provided a detailed view of entorhinal–DG synaptic inputs at specific dendritic compartments and DG granule cell outputs via somatic action potentials (Figure 5A-B). The laminar probe design, with contacts spanning from the pyramidal cell layer of CA1 through the dentate gyrus suprapyramidal blade (GCL1), hilus, and infrapyramidal blade (GCL2), was used to assess circuit throughput (i.e., cortical synaptic input, dendritic compartment integration, and somatic output). Following physiology or behavior a subset of animals were perfused and histology was used to determine the location and percentage of retrovirus infected neurons relative to the laminar probe tract (Figure 5C). We found that the average rate of infection at the site of injection was 18.84±3.37% of *Pten^flx/flx^*and 20.27±3.80% for *Pten^flx/flx^;Raptor^fl/xflx^* (mean±sem, n=15 and 12 mice, respectively).

**Figure 5.**
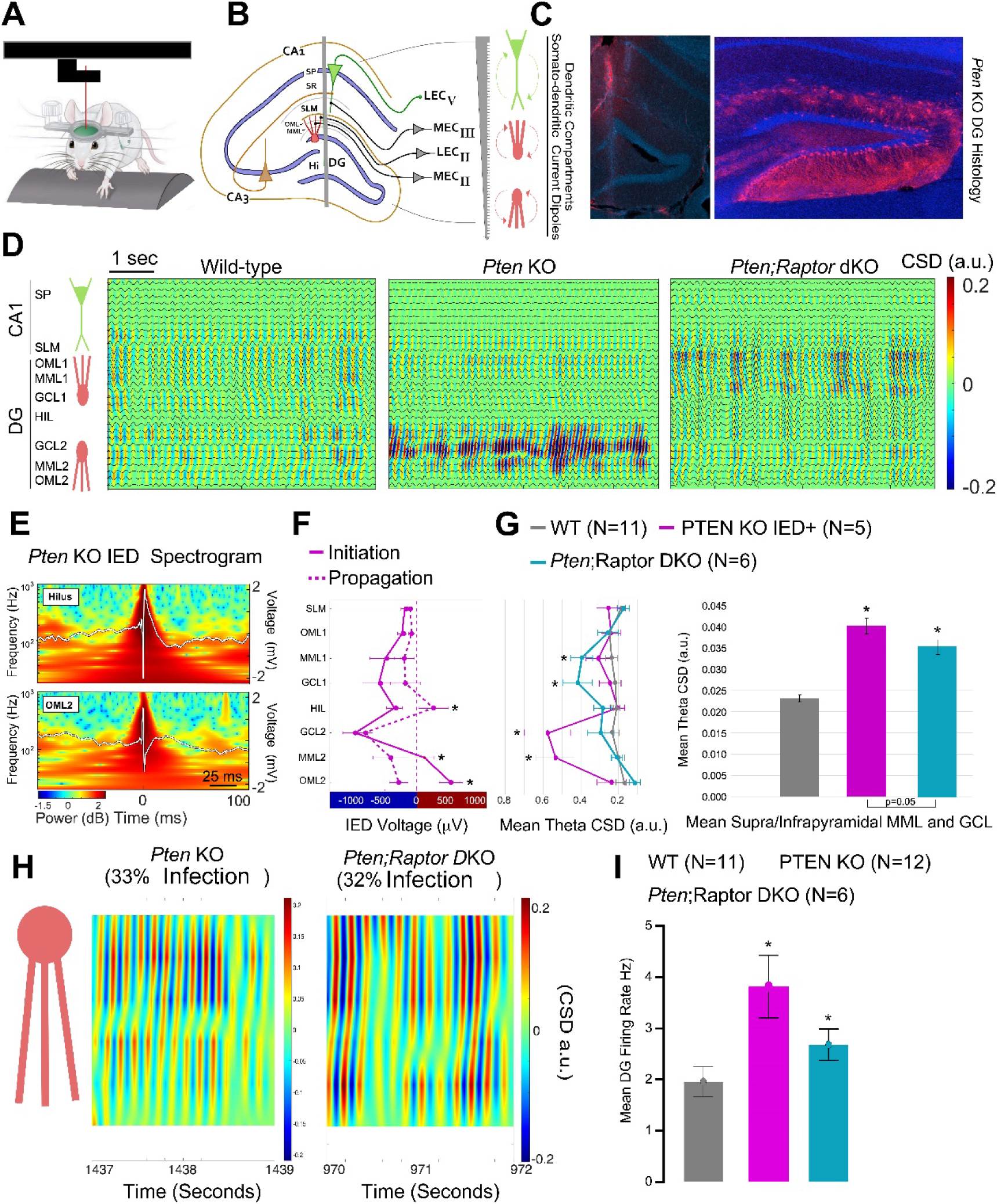
*Pten* and *Pten;Raptor* dKOs Have Distinct Hyperexcitability Signatures *in vivo*. (A) Head-fixed high-density silicon probe recordings from mice during voluntary wheel running. (B) Hippocampal anatomy illustrating electrode position and oscillatory measurements relative to CA1 and dentate gyrus (DG) granule cell layers and their entorhinal cortex projections: CA1 stratum lacunosum moleculare (SLM; layer 3 medial entorhinal cortex, MEC), DG outer molecular layer (OML; layer 2 lateral entorhinal cortex, LEC), and DG middle molecular layer (MML; layer 2 MEC) dendritic compartments. (C) Representative hippocampal histology showing probe track through CA1 and DG (left) and immunohistochemistry illustrating *Pten* knockout cells (tdTomato, red) relative to all cells (DAPI, blue). This example was primarily in the infrapyramidal blade of the DG (right). (D) Examples of current source density (CSD) maps with local field potential (LFP) overlays illustrating depth-resolved theta epochs in wild-type (left), *Pten* KO (middle), and *Pten;Raptor* dKO (right) mice. CSD rasters were rendered using a fixed color scale (±0.2 a.u.) to enable comparison of current sinks (blue) and sources (red) for each subject at each cell layer and along each somatodendritic axis. Theta synchrony is elevated in the upper DG of the *Pten;Raptor* dKO mouse and in the lower DG of the *Pten* KO mouse. (E) Time–frequency scalograms from *Pten* KO from two representative channels (Top = Hilus; Bottom = OML2) with overlaid voltage traces (white) indicating IED timing relative to spectral power changes and high-frequency oscillations. The IEDs represent a fast spike (∼1ms) followed by dendritic propagation. (F) Depth-resolved mean voltage raster profiles comparing *Pten* KO IED initiation (solid red line, −1 to 0 ms) and propagation (dashed red line, 2–15 ms), revealing significant CSD polarity shifts along the GCL2 somatodendritic axis and hilus (GEE, *P* < 0.05). Regions of initiation and propagation on the infrapyramidal blade of DG correlate with higher theta CSD currents. (G) As *Pten;Raptor* dKO do not exhibit IEDs, the primary comparison across groups is theta CSD; Left - Mean theta CSD profiles across groups relative to cell layers and CA1 and DG dendritic compartments (wild-type (gray), *Pten* KO IED+ (mice displaying interictal epileptiform discharges, magenta), *Pten;Raptor* dKO (cyan)). *Pten* KO and *Pten;Raptor* dKO mice exhibit elevated CSD in the granule cell layer (GCL) and MML relative to wild-type (GEE, *P* < 0.05). Right - Mean theta CSD profiles for each group averaged across GCL and MML layers in the infrapyramidal and suprapyramidal blades of the DG wild-type (gray), *Pten* KO IED+(magenta), *Pten;Raptor* dKO (cyan). *Pten* KO and *Pten;Raptor* dKO mice exhibit similar CSD elevation (GLM, P = 0.05), while both groups are significantly larger than wild-type (GLM, *P* < 0.05). (H) Direct comparison between a *Pten* KO and a *Pten;Raptor* dKO mouse exhibiting similar infection rates (∼ 30%) in the infrapyramidal blade of DG, as well as similar spatial CSD levels. (I) Dentate gyrus granule cells in both *Pten* KO (55 cells from N = 12 mice) and *Pten;Raptor* dKO (52 cells from N = 6 mice) exhibit higher mean firing rates than wild-type mice (68 cells from N = 11 mice; GLM, P < 0.05).

Relative to wild-type, both *Pten;Raptor* dKO and *Pten* KO mice exhibited elevated current source density (CSD) levels on the DG somatodendritic axis (Figure 5D). This result indicates hypersynchronous currents relative to entorhinal synaptic inputs and the granule cell layer within the DG. These were highest within a subset of *Pten* KO mice (5/13) that also exhibited interictal epileptiform discharges (IEDs; Mean = 8.2 ± 1.77 events within ∼ 30 min recording sessions). The initiation of the IEDs are anchored at 0 ms, where the example of the corresponding waveforms in Figure 5E denote a positive voltage signal on the somatodendritic axis of GCL2 and negative voltage signal within the hilus and granule cell layers (initiation phase = -1 to 0 ms), followed by a polarity reversal with a positive signal within the hilus and negative signal on the GCL2 axis, representing propagation of the large voltage IED through the DG and hippocampus (propagation phase = 2 to 15 ms). Spectral analysis also indicates these IEDs are associated with high frequency oscillations up to 1000Hz. For this animal, these results correlate with its histology where *Pten* KO was greatest in GCL2 (Figure 5C, Right). This stereotyped pattern of IED waveforms was similar across *Pten* KO animals with IEDs (N=5), where the IEDs originated from the somatodendritic axis below GCL2 (Figure 5F).

Because *Pten;Raptor* dKO mice did not exhibit IEDs, we used theta hypersynchrony via CSD to make direct group comparisons. Using the wild-type hilar region as a comparator (p > 0.05 compared with all other wild-type anatomical regions), we found a significant interaction between group and anatomical region (Generalized Linear Model (GLM), Wald value = 931.41, p < 0.001) (Figure 5G, Left). To account for the possibility of differential virus infection in the infra and suprapyramidal blade, we also conducted an analysis averaging across both granule and molecular layers in both blades (Figure 5G, Right). We also compared the Infrapyramidal somatodendritic axis via CSD for individual *Pten;Raptor* dKO and *Pten* KO mice that the most similar virus infection rates of approximately 30% of DGCs (Figure 5H). Both mice exhibited similar CSD levels across the axis. Finally, we examined the mean DGC firing rate for single cells isolated using kilosort and found that both the *Pten* KO and the *Pten;Raptor* dKO displayed increased firing rates *in vivo* (Figure 5I). Our results indicate that both *Pten* KO and the *Pten;Raptor* dKO have hyperactive dentate gyri as assessed by current source density and mean firing rate. This was more pronounced for the *Pten* KO and interictal epileptiform discharges were only detected in the *Pten* KO.

### *Pten* KO and *Pten;Raptor* dKO Exhibit Differences in Associative Learning and Spatial Navigation Memory

We next examined whether the electrophysiological differences in action potential firing and corticohippocampal circuit efficacy affect spatial learning behavior. Mice are trained to associate approaching an object at a fixed location with receiving a food reward. In phases 1–4, the goal zone around the object became progressively smaller (51 to 15 cm; Figure S3A), while in phases 5–7, the dwell time required within the zone to trigger a reward increased (750 ms to 1.2 s; Figure S3B). The wild-type (n = 19) and *Pten;Raptor* dKO (n = 13) mice learned this task in the same amount of time (p > 0.05). However, *Pten* KO mice (n = 13) required more training trials than wild-type mice to reach the task criterion of 20 rewards within a 30-minute session (GLM; Wald value = 33.594, p < 0.001; Figure 6A). ROC analysis showed that the number of training trials required to reach criterion was significantly predictive of higher bilateral retroviral infection rates, and therefore greater numbers of knockout neurons in both hippocampi, significantly predicted the number of training trials required to reach criterion (AUC > 0.8; Figure 6B). In the first probe session of phase 7, the object was removed, arena cleaned, and reward given for visiting the prior object location. This tested hippocampal-dependent memory for the goal location without the object or scent marking as sensory cues (Figure 6C).

**Figure 6.**
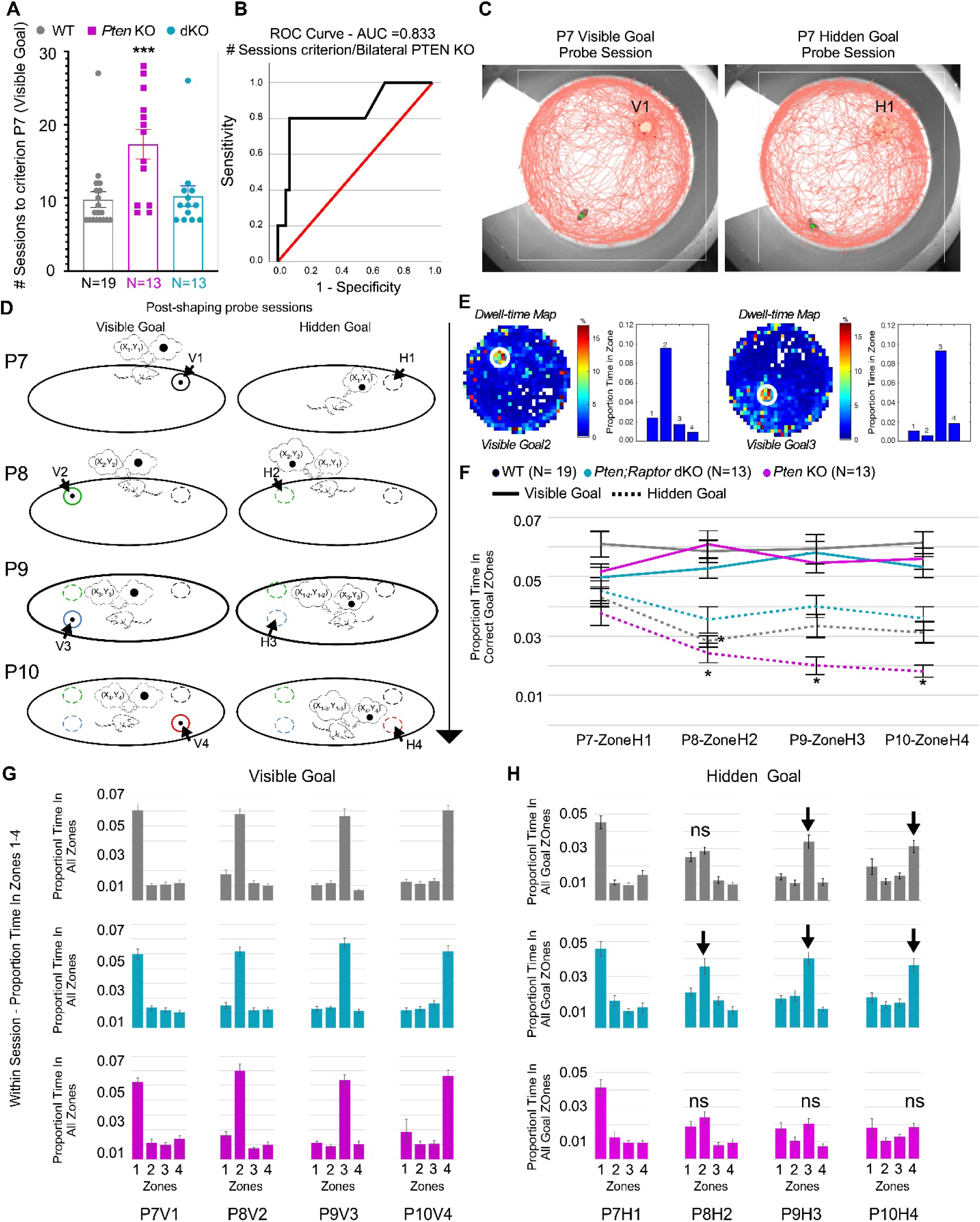
*Pten* KO Mice Have Poor Object and Location Memory while *Pten;Raptor* dKO Mice Have Improved Object-Location Coupling. (A) Number of training sessions required to reach phase 7 and criterion (20 rewards within 30 min) for wild-type (gray), *Pten* KO (magenta), and *Pten;Raptor* dKO (cyan) mice. There was no significant difference between wild-type and *Pten;Raptor* dKO mice (GEE, *P* > 0.05), whereas *Pten* KO mice required significantly more trials than wild-type mice (GEE, *P* < 0.001). (B) Receiver operating characteristic (ROC) analysis demonstrating that bilateral DG Pten deletion rate is a significant predictor of trials to criterion in *Pten* KO mice (area under the curve, AUC > 0.8). (C) Representative performance during visible (V1, left) and hidden (H1, right) goal trials in a 76 cm diameter arena with a polarizing white cue card on a gray background. (D) Following shaping in the primary visible goal position during phase 7 (V1, solid circle), the goal cue was removed and mice were reintroduced to a clean arena for the hidden goal session (H1, dashed circle). Phases 8–10 consisted of paired probe trials conducted on a single day, in which the visible goal was relocated to a different quadrant for one session (V2–V4), followed by a corresponding hidden goal session (H2–H4). This design tests flexible navigation to either the visible goal or its remembered location. (E) Representative spatial accuracy performance from a wild-type mouse during V2 and V3, showing dwell time maps illustrating goal pauses within the goal zone (left) and quantification of the proportion of time spent in the goal zone versus equidistant control zones in the other quadrants (right). (F) Proportion of time spent in the correct goal zone across visible (solid lines) and hidden (dashed lines) sessions for wild-type (gray, N = 19), *Pten* KO (magenta, N = 13), and *Pten;Raptor* dKO (cyan, N = 13) mice. During visible sessions, there was no significant difference within or across groups between baseline and probe sessions (GEE, P > 0.05). During hidden sessions, *Pten;Raptor* dKO were the only group that never differed from H1, while wild-type showed decreased dwell in H2 compared to H1 and *Pten* KO performance declined across zones 1–4 and was significantly lower than *Pten;Raptor* dKO in zones 2–4 (GEE, *P* < 0.05). The average time in the correct goal zone also trended higher in *Pten;Raptor* dKO mice than WT mice across sessions. (G) Within-session proportion of time spent in each zone during visible goal sessions 1–4 and hidden goal sessions 1–4. Wild-type (gray), *Pten;Raptor* dKO (cyan), and *Pten* KO (magenta) tracked the visible object in all locations. (H) Wild-type, *Pten;Raptor* dKO and *Pten* KO performed similarly in hidden goal 1 (Phase 7H1). During phase 8 (H2), only the *Pten;Raptor* dKO mice showed enhanced extinction of the prior H1 location (black arrows). Across probe sessions (H3–H4), *Pten;Raptor* dKO and wild-type mice performed similarly (*P* > 0.05). *Pten* KO mice did not spend more time in the correct goal zone other than the primary H1 goal location. During H3 and H4 *Pten* KO performance was significantly reduced (black arrows) relative to *Pten;Raptor* dKO and wild-type mice (*P* < 0.05). While *Pten;Raptor* dKO demonstrated an enhanced extinction of prior goal zone responses, the *Pten* KO exhibited a perseverative response on the primary goal zone.

Mice were next subjected to a series of trials in which the object was first placed in its training location, and mice approached and remained within the goal zone for 1.2 s to receive a reward (i.e., visible location 1, V1). The object was then removed, arena cleaned, and mice were required to approach the previous object location to receive a reward (hidden location 1, H1). On subsequent days, the object was relocated to new quadrants (V2–V4), each followed by a hidden trial (H2–H4) (Figure 6D). Examples of performance by visible goal rotation are shown in Figure 6E. Across probe sessions, generalized estimating equation (GEE) analysis revealed a significant Group × Training Phase interaction for the proportion of time spent in the goal zone during hidden goal probe sessions (Wald χ² = 104.57, df = 11, p < 0.001), with WT at H1 serving as the statistical comparator. *Pten* KO mice displayed a progressive and significant decrease in goal-zone occupancy across sessions H2–H4 relative to WT baseline (H2: Wald χ² = 14.46, p < 0.001; H3: Wald χ² = 21.11, p < 0.001; H4: Wald χ² = 43.27, p < 0.001), indicating a failure to associate the object with new goal locations across sessions. In contrast, *Pten;Raptor* dKO mice were the only group that did not differ significantly from WT baseline at any post-baseline session (H2: Wald χ² = 1.74, p = .187; H3: Wald χ² = 0.36, p = .551; H4: Wald χ² = 1.80, p = .180), demonstrating preserved capacity to associate the object with successive novel goal locations. WT control mice showed the expected pattern of increased goal-zone occupancy across sessions (H2: p < 0.001; H4: p = .012), consistent with normal extinction of prior spatial associations. Taken together, these findings indicate that *Pten* KO mice fail to flexibly update object–location associations across successive probe sessions, whereas *Pten;Raptor* dKO mice retain this capacity. The behavioral differences observed in *Pten* KO mice were not attributable to anxiety-related behavior or thigmotaxis, as no group differences were found in the proportion of time spent in the center (Wald value = 6.58, p = 0.832) or annulus of the cylinder (Wald value = 23.47, p = 0.15) (Figure S4).

Within-session analysis of the proportion of time spent in zones 1–4 of each quadrant revealed significant Group × Training Phase × Zone interactions for visible goal sessions (GEE; Wald value = 2.98 × 10¹⁵, p < 0.001) and hidden goal sessions (GEE; Wald value = 6789.24, p < 0.001). Despite group differences across visible goal sessions, within-session analysis showed that mice in all three groups spent significantly more time in the visible goal zone than in any of the other three zones (Figure 6G). Within-session analysis of hidden goal sessions showed similar performance between wild-type and *Pten;Raptor* dKO mice across probe sessions, whereas *Pten* KO mice perseverated on zone H1 across goal rotation sessions (Figure 6H; Figure S5). In the first goal rotation session (H2), group differences were also observed in extinction of unrewarded pauses in the first goal zone (H1). Both wild-type and *Pten* KO mice spent similar amounts of time in H1 and H2 (p > 0.05), whereas *Pten;Raptor* dKO mice spent significantly more time in the new goal zone during phase 8 (H1 versus H2; p = 0.01).

### *Pten;Raptor* x POMC-Cre dKO Have Non-Convulsive Seizures

Retroviral infection at P7 resulted in Cre expression in 19.48±2.48% of dentate granule neurons at the site of injection across all animals. None of these animals displayed overt behavioral seizures. In 60-day old POMC-Cre+ x Ai14 x Pten^flx/flx^ and *Pten^flx/flx^;Raptor^flx/flx^* mice tdTomato was expressed in 83.0 ± 3.3% of PROX1 expressing dorsal granule neurons (Figure 7A; n=4 *Pten* KO and 3 *Pten;Raptor* dKO mice). Qualitatively, the *Pten* KO dentate gyrus exhibited foliation, which was not observed in Cre- or *Pten;Raptor* dKO mice (Figure 7B). Using acute hippocampal slice electrophysiology, we found that dentate granule neurons in these animals exhibited significant burst firing and altered fast AHP, phenocopying the effects observed in prior retroviral conditional knockout experiments (Figure S6).

**Figure 7.**
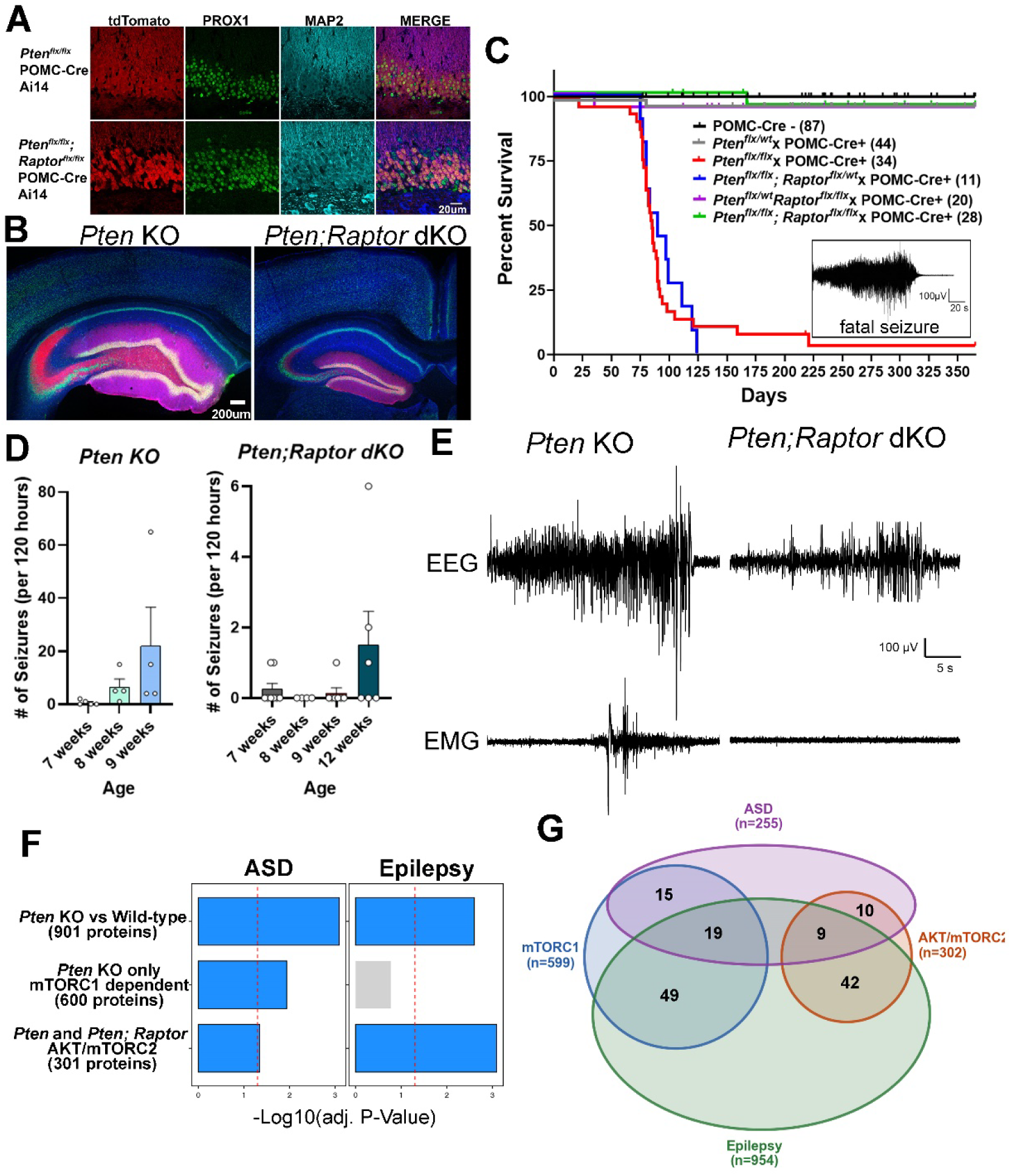
*Pten^flx/flx^*× POMC-Cre and *Pten^flx/flx^;Raptor^flx/flx^* × POMC-Cre Have Seizures and Share a Molecular Signature with ASD and Epilepsy. (A) Representative hippocampal histology from *Pten^flx/flx^* × POMC-Cre+ × Ai14 and *Pten^flx/flx^;Raptor^flx/flx^* × POMC-Cre+ × Ai14 mice stained for tdTomato (red), granule neurons (PROX1, green), and dendrites (MAP2, cyan), demonstrating the percentage of Cre+ cells and basal dendrite sprouting into the dentate hilus of the *Pten* KO. (B) Low-magnification hippocampal sections stained with NeuN (green), MAP2 (blue), and tdTomato (red). The *Pten^flx/flx^* × POMC-Cre+ granule cell layer (GCL) shows pronounced foliation consistent with epileptic pathology, whereas *Pten^flx/flx^;Raptor^flx/flx^* × POMC-Cre+ mice do not. (C) Kaplan–Meier survival curve showing near-complete mortality of *Pten* KO mice (red) between 8–10 weeks of age; inset, representative EEG recording during a fatal seizure in a Pten KO mouse. Mortality is rescued in *Pten;Raptor* dKO mice (green). The POMC-Cre-, Pten^flx/wt^ × POMC-Cre+ and *Pten^flx/flx^* × POMC-Cre+ control groups have been previously published (Prina et al., 2026) (D) Continuous video-EEG monitoring of *Pten^flx/flx^* × POMC-Cre+ and *Pten^flx/flx^;Raptor^flx/flx^* × POMC-Cre+ mice demonstrating seizure onset at 7–8 weeks in *Pten* KO animals, with all individuals exhibiting frequent seizures by 9 weeks. *Pten;Raptor* dKO mice display electrographic non-convulsive seizures. (E) Representative EEG/EMG traces illustrating a generalized tonic–clonic seizure in a *Pten* KO mouse and a non-convulsive seizure, with no EMG activity elevation typical of *Pten;Raptor* dKO mice. (F) Hypergeometric enrichment analysis showing overlap between differentially regulated proteins and disease-associated gene sets for autism spectrum disorder (ASD), and epilepsy. Red dashed line indicates adjusted significance threshold. (G) Four-way Venn diagram showing the overlap between PTEN-regulated phosphoproteins (mTORC1-dependent, n = 600, blue; AKT/mTORC2-dependent, n = 301, orange) and curated human disease gene lists for ASD (Fu et al., 2022; FDR ≤ 0.1; n = 255, purple) and monogenic epilepsy (Genes4Epilepsy v2023-03, Oliver et al., 2023; n = 954, green). Numbers within each intersection indicate the count of genes shared across the indicated sets. The triple-overlap regions (19 and 9) identify genes shared across the PTEN phosphoproteome, ASD, and monogenic epilepsy.

Consistent with previous studies, increasing the number of knockout cells in the dentate gyrus beyond a threshold resulted in the development of overt seizures (LaSarge et al., 2021; Pun et al., 2012; Santos et al., 2017). *Pten^flx/flx^* x POMC-Cre+ animals began exhibiting generalized tonic–clonic seizures in video-EEG-EMG recordings at 7 weeks of age, with seizure frequency increasing over time (4.4 ± 2.9 seizures per day at 9 weeks of age), followed by lethality by 10 weeks (Figure 7C-E). One animal was captured on video dying during a SUDEP event (inset, Figure 7C), suggesting that terminal seizures contribute to the early lethality phenotype. By contrast, *Pten^flx/flx^;Raptor^flx/flx^* x POMC-Cre+ animals, showed a complete rescue of mortality, surviving for over one year (Figure 7C). Video-EEG-EMG analysis of *Pten^flx/flx^;Raptor^flx/flx^* x POMC-Cre+ animals also showed a rescue of the generalized tonic-clonic seizure phenotype, as no animals (0/9) experienced a generalized tonic–clonic seizure.

However, five of nine *Pten^flx/flx^;Raptor^flx/flx^*x POMC-Cre+ animals displayed electrographic seizures in the EEG without a corresponding change in the EMG (0.2 ± 0.1 per day at 12 weeks of age (Figure 7D,E). Together with the whole-cell electrophysiology results, these data suggest that *Raptor*-dependent morphological and synaptic changes are necessary for the most severe consequences of *Pten* loss, whereas the *Rictor*-dependent burst-firing phenotype remains sufficient to drive a lower level of circuit hyperexcitability.

### mTORC1 and AKT-mTORC2 Regulated Proteins Overlap with ASD, and Epilepsy Genes

To determine whether our phosphoproteomic data supported a molecular signature consistent with human disease, we compared the 901 PTEN-regulated phosphoproteins with curated gene lists from ASD and epilepsy. Hypergeometric enrichment analysis revealed significant overlap with both disease categories (Figure 7F). Both the mTORC1-dependent and AKT/mTORC2-dependent protein sets were enriched for ASD-associated genes ((Fu et al., 2022); FDR ≤ 0.1; n = 255), consistent with the broad contribution of PTEN signaling to ASD risk. In contrast, the overlap with monogenic epilepsy genes (Genes4Epilepsy; (Oliver et al., 2023); n = 954) was proportionally greater in the AKT/mTORC2-dependent set (51/302, 16.9%) than in the mTORC1-dependent set (68/599, 11.4%), directly paralleling the growth versus excitability partition we identified at the molecular level. We identified 53 of 255 ASD-risk genes from Fu et al. 2022, and 119 of 954 epilepsy-risk genes from Oliver et al. 2023 within the 901 PTEN-regulated phosphoproteins (Figure 7G, Supplemental Tables).

Nearly all of the epilepsy-associated phosphoproteins across both arms are implicated in developmental and epileptic encephalopathy (DEE), the most severe and pharmacoresistant end of the epilepsy spectrum, suggesting that PTEN-regulated phosphorylation converges on the molecular substrates of highly penetrant seizure disorders. Strikingly, 28 of the 53 ASD-risk phosphoproteins (52.8%) were also present in the epilepsy gene list, identifying a core set of high-confidence candidates shared across the PTEN phosphoproteome, ASD, and epilepsy simultaneously. Within the mTORC1-dependent triple-overlap set (19 genes), *KCNMA1*, encoding the BK channel and among the most significant ASD-risk genes in this group (FDR = 0.0003). The remaining mTORC1 triple-overlap genes were enriched for chromatin regulators and transcriptional scaffolds, consistent with mTORC1’s established role in translational control. Within the AKT/mTORC2-dependent triple-overlap (9 genes) were the postsynaptic scaffolding proteins *SHANK3* and *SYNGAP1*, the calcium/calmodulin-dependent kinase *CAMK2A*, the voltage-gated calcium channel *CACNA1E*, and the sodium channel *SCN2A*, further linking the AKT/mTORC2-driven excitability program to the molecular substrates of ASD-associated seizure susceptibility. The functional contrast between the two triple-overlap sets, mTORC1 enriched for chromatin factors and *KCNMA1* and AKT/mTORC2 dominated by postsynaptic scaffolds and ion channel subunits, mirrors the electrophysiological bifurcation we characterized and suggests that the two arms of PTEN signaling contribute to overlapping but mechanistically distinct dimensions of ASD and epilepsy risk.

## Discussion

Neurons must coordinate their growth and intrinsic excitability during development to maintain stable output as their morphology changes (Marder and Goaillard, 2006; Turrigiano and Nelson, 2004). Neuronal growth increases membrane surface area and, with it, total membrane capacitance. Because membrane capacitance scales with area, a neuron that simply adds ion channels in proportion to its growth would require increasingly large absolute currents to achieve equivalent membrane depolarization, predicting reduced intrinsic excitability per unit of synaptic input despite greater absolute ionic current. One mechanism by which PTEN-deficient neurons compensate for this size-excitability paradox is a disproportionate expansion of excitatory synapse number, which augments total synaptic drive onto enlarged cells (Williams et al., 2015). Canonically, growth factors increase downstream MAPK, PLCγ, PI3K and mTORC1/2 signaling in a coordinated fashion (Kaplan and Miller, 2000). While there is cross-talk among signaling cascades, MAPK largely influences transcription, mTORC1 translation and lipid synthesis. Here, we demonstrate active scaling of intrinsic excitability through mTORC2-dependent phosphoregulation of voltage-gated Ca^2+^ and BK channels, providing a cell-autonomous means of calibrating membrane excitability to match the growth state of the neuron.

The AKT-mTORC2 program controls a separable set of cellular phenotypes. Genetic deletion of *Akt* or the obligatory mTORC2 gene *Rictor* in *Pten* knockout neurons rescues the decreased fAHP and burst firing without rescuing growth or synaptic overgrowth. Electrophysiological dissection in *Pten;Raptor* dKO neurons identified reductions in voltage-gated K^+^ and Ca^2+^ currents that underlie the altered firing. Selective blockade of individual K^+^ channels indicated a disproportionate loss of BK current, and *Kcnma1* deletion recapitulated the fAHP reduction. Ca^2+^ current reductions affected P/Q, N, and R-type subtypes. The reduction in aggregate Ca^2+^ current accompanying increased burst firing is initially counterintuitive, because Ca^2+^ entry depolarizes the membrane and would be expected to support firing (O’Leary et al., 2014). We hypothesize that the spatial coupling of CaV channels to BK channels may be disrupted. BK channel activation requires Ca^2+^ entry through nearby voltage-gated Ca^2+^ channels producing nanodomains delivering high local Ca^2+^ to BK channels during the action potential (Berkefeld et al., 2006; Fakler and Adelman, 2008; Indriati et al., 2013). This coupling underlies the fAHP, which limits the rate of repetitive firing. Reduced somatic CaV current, particularly if accompanied by altered CaV-BK colocalization, would weaken BK activation, attenuate the fAHP, and release the brake on repetitive firing. Phosphoproteomic analysis revealed altered phosphorylation of voltage-gated calcium and potassium channel subunits in *Pten;Raptor* dKO neurons, providing a substrate for AKT-mTORC2-dependent regulation of channel function, trafficking, scaffolding, or subcellular targeting. Whether the relevant change is altered single-channel conductance, altered surface expression, altered subcellular localization, or altered scaffolding interactions remains to be resolved.

Behavioral and in vivo physiological analyses reveal both mTORC1-dependent and mTORC1-independent phenotypes. Dentate gyrus specific *Pten* knockout produces impairments in object and spatial memory (Getz et al., 2022). *Raptor* co-deletion rescues the principal *Pten* behavioral phenotype, difficulty pairing object with reward and inability to flexibly associate new locations with the object. *Pten;Raptor* dKO mice were better able to associate an object with new goal locations raising the possibility that increased excitability with normal synaptic inputs enhances dentate gyrus pattern separation (Anacker and Hen, 2017; Anacker et al., 2018; Garthe et al., 2009). Further, bursting dentate granule neurons in *Bmal1* KO or with optogenetic activation can improve object memory (Ewell et al., 2019; Gonzalez et al., 2023; Gonzalez et al., 2025). Although the behavioral phenotype in *Pten;Raptor* dKO mice is subtle, the increased sensitivity of *in vivo* electrophysiology reveals a clear signature of network hyperactivity even when growth has been rescued.

Clinical mutations in ASD-associated genes rarely produce ASD with complete penetrance. Estimates of ASD penetrance among individuals with SCN2A mutations range from 17–50% (Kruth et al., 2020; Richardson et al., 2022; Sanders et al., 2018), while penetrance associated with SYNGAP1 mutations may be as high as 68% (Wiltrout et al., 2024). In PTEN hamartoma tumor syndrome, ASD penetrance is approximately 25% across a meta-analysis of 14 cohorts (Cummings et al., 2022). Among the 255 genes in which de novo mutations have been identified in individuals with ASD at FDR ≤ 0.1 (Fu et al., 2022), 53 encode proteins whose phosphorylation is altered downstream of PTEN. We hypothesize that gene-gene interactions within this network influence whether individuals with PTEN hamartoma tumor syndrome meet diagnostic criteria for ASD, and that similar interactions may account for a substantial fraction of ASD cases more broadly. We found a clear overlap of the phophoproteomic signature of *Pten* KO neurons both downstream of mTORC1 and mTORC2 and ASD. This broad overlap may be due to the heterogeneity of ASD.

Genetic or pharmacological suppression of mTORC1 rescues neuronal hypertrophy, dendritic and spine overgrowth, and many associated behaviors in mouse models of PTEN loss and related disorders (DeSpenza et al., 2025; Getz et al., 2016; Godale et al., 2022; Godale et al., 2026; Narvaiz et al., 2023; Tariq et al., 2022; Zhou et al., 2009). This has motivated clinical trials targeting mTORC1 to treat cognitive and behavioral symptoms in patients with PTEN mutations and ASD (Srivastava et al., 2022). Improvements were significant but incomplete, raising the possibility that either post-developmental treatment is inadequate or that mTORC1-independent mechanisms contribute to symptoms. Our results support the latter possibility directly. *Raptor* co-deletion in *Pten* knockout neurons rescues cellular size and excitatory synaptic input, yet leaves intrinsic hyperexcitability and non-convulsive electrographic activity intact. The *Pten;Raptor* x POMC-Cre mice fully rescue mortality and tonic-clonic seizures, yet electrographic abnormalities and non-convulsive seizures persist. Dendritic overgrowth is problematic for hippocampal networks in both acquired and genetic models (Kumari and Brewster, 2024) and may be irreversible after development. By contrast, ion channel dysfunction may be more amenable to pharmacological intervention to offset network hyperactivity and altered spike timing.

Our findings generate specific testable predictions at both the cellular and circuit levels. First, the apparent paradox of reduced aggregate Ca^2+^ current accompanying increased burst firing implies that channel localization and coupling, rather than total channel current, determine excitability. Two-photon Ca^2+^ imaging or voltage imaging in somatic and dendritic compartments would directly test whether somatic CaV-BK nanodomains are weakened while dendritic Ca^2+^ signaling is preserved or amplified. Second, our behavioral findings raise specific questions about how altered intrinsic excitability shapes hippocampal coding during learning. Recording place cells in dentate granule, CA1, and CA3 neurons in freely moving *Pten* and *Pten;Raptor* dKO mice during the spatial accuracy task would reveal whether the behavioral phenotype reflects altered place field formation, altered place field stability across sessions, or altered pattern separation between similar contexts. Combining this with recordings of object cells in lateral entorhinal cortex and place cells in medial entorhinal cortex would test our prediction that the proximal-distal dendritic shift in *Pten;Raptor* dKO neurons biases the hippocampus toward MEC-driven spatial coding at the expense of LEC-driven object coding. Overall, we demonstrate a novel mechanism through which mTORC2 regulates excitability. This mechanism impacts physiological and behavioral output using the mouse dentate gyrus as a test circuit. Finally, the phosphoproteomic signature of mechanisms regulating neuronal growth and activity in mice show a clear overlap with high-confidence genes associated with epilepsy and ASD.

## ACKNOWLEDGEMENTS

We would like to thank Heinz Beck, Farah Lubin, Jacques and Linda Wadiche for valuable feedback on this work. This work was supported by the Dr. Paul R. Jarvis HOPE Endowment for Epilepsy (JMB), UAB Office of Research and UAB’s Civitan International Research Center, a multi-investigator grant from the PTEN Research Foundation, a charity governed by English law (charity number 117358) to BWL, JMB, and MCW under grant number UAB-24-001, NIGMS T32GM146611 (MLP), NIMH R01 MH097949 (BWL), NIMH R01 MH134787 (JMB), NINDS R21 NS133670 (JMB), NINDS R01NS110945 (MCW), and NINDS R01NS111029 (KTK). The proteomics experiments were done as part of an internship awarded to AFA by IDeA National Resource for Quantitative Proteomics under NIH grant R24GM137786. Thanks to Isabella Piatek for behavioral training and manual scoring of IEDs, Caitlyn Marczely and Eli Rachimi for behavioral training.

## AUTHOR CONTRIBUTIONS

Study design and conceptualization: BWL, JMB, MCW

Slice Electrophysiology and Analysis: AFA, BWL

Stereotaxic Virus Injections: AFA, MLP

Histology and Image Analysis: AFA, MLP

In Vivo Electrophysiology and Analysis: JMB, NE, RY, HM, MB

Mouse Behavior and Analysis: JMB, DM, NE, HH

Video EEG/EMG Monitoring and Analysis: MCW, JU

Phosphoproteomics: AFA, KEB, DWP

Bioinformatic Analysis: ED, AFA, BWL, KTK

Writing and review of manuscript: BWL, JMB, MCW, KTK, AFA, ED

Funding acquisition and supervision: BWL, JMB, MCW, KTK

## Declaration of generative AI and AI-assisted technologies

ChatGPT and Claude Code were used to generate novel python scripts for data analysis and to edit writing clarity and ensure no grammatical errors in the manuscript. After using this tool/service, the authors reviewed and edited the content as needed and take full responsibility for the content of the publication.

## Methods

### Animals

All animal procedures were conducted in accordance with the National Institutes of Health Guide for the Care and Use of Laboratory Animals and were approved by the Institutional Animal Care and Use Committee (IACUC) at respective institutions: Luikart Animal Protocol Number (APN): IACUC-23042, Weston (IACUC- 25-186 (FBRI)), and Barry (PROTO202000145). Mice were housed in a temperature- and humidity-controlled vivarium on a 12 h light/dark cycle with ad libitum access to food and water. Every effort was made to minimize animal suffering and to reduce the number of animals used.

### Stereotaxic Injections

Postnatal day 7 (P7) mouse pups were anesthetized with 4% isoflurane and secured in a nose cone through which isoflurane was continuously delivered. Body temperature was maintained using a heating pad. The scalp was treated topically with betadine and lidocaine before a midline incision was made. Isoflurane concentration was then reduced to 1.5–2%, and surgical coordinates were set relative to lambda. Using a 10 µL Hamilton syringe, up to 2 µL of suspended retroviral solution was injected bilaterally into the dentate gyrus at a rate of 0.3 µL/min at the following coordinates relative to lambda: x = ±1.3 mm, y = +1.55 mm, and z = −2.3, −2.2, −2.1, and −2.0 mm. At each z-depth, 25% of the total volume was delivered. After completion of the injections, the syringe was held at z = −2.0 mm for 2 min, slowly withdrawn to z = −1.0 mm, held for 1 min, and then slowly removed from the brain. Following bilateral injections, the scalp incision was sutured and a topical antibiotic was applied. Pups were ear-marked for identification and placed in a heated recovery chamber until fully ambulatory.

### Histology & Image Quantification

For all histological analysis, mice were transcardially perfused with ice-cold 1x PBS-Sucrose (4%) and fixed with 4% paraformaldehyde (PFA). Brains were post-fixed for 24 hours in 4% PFA and 50µm thick vibratome sections made (LeicaVT1200S). To determine the percent of granule neurons infected in animals that had undergone behavior and *in vivo* electrophysiology, slices were then immunostained for Prox1 to visualize the granule neurons. For control animals injected with retroviruses expressing GFP (no Cre) they were co-stained for GFP. For *Pten*^flx/flx^ x Ai14 knockout animals in which retroviral Cre was expressed, anti-RFP was used to visualize tdTomato to quantify the percentage of Cre expressing neurons. For *Pten*^flx/flx^ ; *Raptor^flx/flx^* animals that Ai14 allele was not present so we used anti-loPten IHC to estimate the percentage of cells expressing Retroviral cre. Freely floating slices were incubated in primary antibody solution for 24-48 hours at 4°C, and then stained with corresponding secondary antibodies from JacksonImmuno Research for 24-72 hours at 4°C. Stained slices were then mounted with DAPI containing Vectashield (Vector Laboratories) and z-stack images were acquired with confocal microscopy (LSM800) at 5X, 10X, and/or 40X magnification with 0.7X zoom, as specified in the figure legends. For animals having undergone *in vivo* electrophysiology we visually located the electrode tract and quantified neurons from adjacent tissue. The percentage of infected neurons was quantified in ImageJ/FIJI using the Multi-point tool, to count all Prox1 labeled granule neurons, and then to count all Cre-expressing neurons within that image plane, and from there a percentage was generated. For the *Pten^flx/flx^,* and *Pten*^flx/flx^, *Raptor^flx/flx^* x Ai14 x POMC-Cre animals were perfused at P60, sectioned and immunostained for RFP (tdTomato), MAP2, Prox1 to determine what percentage of the granule neurons express POMC-Cre.

| Antibody | Manufacturer, Catalog# | Dilution |
| --- | --- | --- |
| Chicken anti-GFP | Abcam, #13970 | 1:3000 |
| Rabbit anti-mCherry | Abcam, #167453 | 1:3000 |
| Mouse anti-RFP | Rockland, #200-301-379 | 1:3000 |
| Mouse anti-NeuN | EMD Millipore, #377 | 1:500 |
| Rabbit anti-Prox1 | Abcam, #199359 | 1:500 |
| Chicken anti-MAP2 | Abcam, #5392 | 1:1000 |
| Rabbit anti-PTEN (138G6) | Cell Signaling Technology #9559 | 1:70 |

### Whole-cell acute slice electrophysiology

Tangential 290uM thick hippocampal slices were generated using an ice-cold cutting solution (all in mM) of 110 CholineCl, 7 MgCl_2_, 2.5 KCl, 1.25 NaH_2_PO_4_*2H_2_O, 0.5 CaCl_2_, 10 dextrose, 1.3 Na-Ascorbate, and 25 NaHCO_3_. Mice are anesthetized with 2% Avertin and are perfused with cutting solution prior to brain dissection. Slices were moved to recording solution 125 NaCl, 25 NaHCO_3_, 1.25 NaH_2_PO_4_*2H_2_O, 2.5KCl, 25 dextrose, 2 CaCl2, 1 MgCl2 and adjusted to ∼290mOsm. All solutions are bubbled with 95% O_2_. Slices were stored at 34°C for 30 minutes and maintained at room temperature. All recordings were performed at ∼37°C using TC-324B (Warner Instruments). Whole-cell current clamp experiments were performed using potassium gluconate internal solution (all in mM) 115 Kgluconate, 10 HEPES, 2 EGTA, 20KCL, 2 MgATP. 10 NaPhosphocreatine, 0.3Na_3_GTP and adjusted to ∼ 290mOsm. Pipettes (PG10165-4, WPI) were pulled using a PC-10 (Narishige) puller to a typical tip resistance of 2-6MΩ. Recordings were acquired using a Multiclamp 700B and Multiclamp Commander (Molecular Devices) and digitized with USB-6221 (National Instruments). R_m_, R_s_, and C_m_ were all calculated after going whole-cell on an average of 20 traces of a 40ms, 10mV test pulse with no whole-cell correction. Test pulses were used throughout experiments to monitor for stabile series resistance. Recording was performed with AxoGraphX (John Clements) using 80Khz sample rates. Voltage clamp experiments were performed using a V_h_ of -90mV, pipette capacitance neutralization, and whole-cell correction with typical parameters of ∼7-10pF at 5-15MΩ. Current clamp experiments used pipette capacitance neutralization of 3 to 4 pF auto bridge balance adjustments typically between 15 to 35MΩ. Action potentials were captured using template matching to a template with a 77.799mV amplitude, 0.4250ms 10-90%rise, and 0.826708ms half-width with a 1ms baseline and a 4ms length with a minimum separation of 2ms a captured baseline of 10ms with a 40ms length and a threshold of 1. Any action potentials with an amplitude of less than 20mV are rejected. Action potential shape was measured for the first action potential at the minimum current step eliciting firing (rheobase) and for the average of every action potential fired at rheobase. Burst probability was defined as the number of action potentials fired with inter-spike intervals of less than 20ms divided by the sum of number of action potential with inter-spike intervals of less than 20ms and greater than 20 at rheobase.

Voltage clamp experiments for potassium currents (*I_K_*) were performed with K gluconate internal solution and 1µM TTX added to the external to block Na+ currents and delivered 50ms steps between membrane potentials of -50mV and +20mV in 10mV increments (Fig4.a). We then washed on inhibitors and analyzed the change in peak current amplitudes. Voltage clamp experiments for calcium currents (*I_Ca_*) were done with Cs gluconate internal solution of the following concentration (mM); Cs Gluconate (113), HEPES (10), EGTA (10), CsCl (17.5), NaCl (8), MgATP (2), NaGTP (0.3), and biocytin at 0.2%, and the external solution was supplied with 1µM TTX and 25mM TEA. The concentration and specification of each channel blocker are below.

| Drug | manufacturer | Target channel | Concentration in μM |
| --- | --- | --- | --- |
| Tetrodotoxin (TTX) | Tocris/ CAS #: 4368-28-9 | Na<sub>v</sub> blocker | 1 |
| Tetraethylammonium (TEA) | Sigma Aldrich/ CAS #:56-34-8 | K<sub>v</sub> blocker | 25 (mM) |
| Paxilline | Alomone labs/ Lot #: P450SM0925 | BK potassium channel blocker | 3 |
| Nifedipine | EMD Millipore/ Lot #: 4164536 | L-type Ca<sub>v</sub> blocker | 10 |
| Z944 | Tocris/ CAS #: 1199236-64-0 | T-type Ca<sub>v</sub> blocker | 10 |
| TTA-P2 | Alomone labs/ Lot #: T155SM0410 | T-type Ca<sub>v</sub> blocker | 5 |
| ω-Conotoxin MVIIC | Alomone labs/ Lot #: C150CN3505 | P/Q and N-type Ca<sub>v</sub> blocker | 1 |
| SNX-482 | Alomone labs/ Lot #: RTS500SX3550 | R-type Ca<sub>v</sub> blocker | 0.5 |
| ω- Agatoxin TK | Alomone labs/ Lot #: STA530TX101 | P/Q-type Ca<sub>v</sub> blocker | 0.2 |
| ω-Conotoxin GVIA | Alomone labs/ Lot #: C300CG5525 | N-type Ca<sub>v</sub> blocker | 1 |
| ω-Conotoxin MVIIA | Alomone labs/ Lot #: C670CN2301 | N-type Ca<sub>v</sub> blocker | 0.05 |
| ML-252(hydrochloride) | Cayman Chemicals/ Batch: 0649667-3 | K<sub>v</sub>7 Blocker | 10 |
| XE991(dihydrochloride) | Alomone lab/ Lot #: X101SM0110 | K<sub>v</sub>7 Blocker | 100 |
| GxtX (Guangxitoxin-1E) | Alomone labs/ Lot #: STG200TX0705 | K<sub>v</sub>2 Blocker | 0.1 |
| α-DTX (α-Dendrotoxin) | Alomone labs/ Lot #: D350DE25014 | K<sub>v</sub>1 Blocker | 0.1 |

Drug sensitivity at each voltage was computed using a custom Python script (DrugSensitivityAtEachVoltage.py), which is publicly available at https://github.com/luikartlabgpt-stack/DrugSensitivityAtEachVoltage

#### Spatial accuracy Behavior

The goal of spatial accuracy training was to shape food restricted mice (85% of baseline body weight) to associate a visible bottle cap cue, placed roughly in the middle of one of the arena quadrants, with automated food reward. As training progressed, the shaping procedure reduced the size of the reward zone, and the amount of time required to trigger a pellet release from an overhead feeder would increase. The goal cue would then be removed to test the animal’s ability to flexibly associate the object with both the original training location and novel spatial locations during probe trials (Bures et al., 1997; Kubie et al., 2007; Mouchati et al., 2020).

Over the course of training, the goal zone diameter ranged down to 10% of the 76 cm diameter arena. The task has two modes for spatial behavior. One mode is for goal-directed navigation when the animal is seeking the goal zone and the other for foraging for a food reward after the pellet has been dropped into a random arena location following a successful pause within the goal zone. The visible and hidden versions of the spatial accuracy task were modeled after the cue and place navigation tasks in the Morris water maze (Morris et al., 1982). Hippocampal lesions cause a significant and long-lasting place navigational impairment that cannot be attributed to motor, motivational or reinforcement deficits. In contrast to the water maze task, the spatial accuracy task allows for continuous measurement of multiple goal navigation epochs and spatial memory measurements in one session (Getz et al., 2022; Mouchati et al., 2020). Mouse location was tracked and recorded via a firewire camera (30 Hz sampling rate) placed over the arena and analyzed with Biosignal software (Tracker, Bio-signal Group Corp, Brooklyn, USA) and custom software (MATLAB v R2023B, MathWorks, Natick, MA). Training began with a 2 cm diameter white bottle cap used as a visible goal. The bottle cap was placed ¾ from the arena wall to the arena center in the northeast quadrant. A polarizing cue card (color code gray 9.5; Color-Aid Corp., Hudson Falls, NY) was placed along the north sector of the arena wall, covering approximately 45⁰ of arc. Animals were advanced to successive visible goal phases after reaching a criterion of 20 rewards. In phases 1-4 of training, 500 ms dwell-time in a target zone successively ranging in diameter from 51 cm (Phase 1), 28 cm (Phase 2), 19 cm (Phase 3) and 15 cm (Phase 4) elicited a +5V TTL pulse via a peripheral component interconnect. This pulse triggered release of a food pellet reward (Bioserv, New Jersey; 20 mg dustless precision pellets) from a custom overhead feeder, which fell to a random arena location. A refractory reward period of 5 sec was set to encourage the animal to leave the target area and forage for the fallen pellet before returning to the goal zone to trigger another pellet. In this manner, measures of continuous navigation to and from the goal zone were possible over the course of each 30-minute training session. In **phases 5-7**, the threshold for target dwell-time was raised to 750 ms, 1 sec, and 1.2 sec consecutively. Finally, in **phase 8,** we conducted hidden goal training. Possible odor cues were controlled for by rotating the floor 45° and cleaning it with soap and water, and then 70% ethanol. The mouse was then placed in the arena, where its pauses near the goal provided a proxy measure of self-localization relative to the goal zone, defined by stationary spatial cues in the room (i.e. cue card on the wall of training room). After reaching the 20-goal criterion during several sessions in the original goal location, pairs of probe test sessions were carried out in each of the 3 remaining quadrants: one with the visible goal cue and then another without the goal cue **(Phases 9-11)**. Only 1 pair of visible/hidden goal probe trials were carried out per day and the mouse only triggered food reward in the newly rotated goal location, at the same distance from the arena wall as the original training location. These probe sessions were typically separated by approximately 10 minutes while the experimenter cleaned and rotated the floor to control for possible odor cues.

#### *In vivo* Electrophysiology

Electrophysiological Recordings and Experimental Logic: Following assessment of spatial cognition on the spatial accuracy task, wideband LFPs (1-6000Hz; sampling rate = 30.3 kHz) were recorded from CA1 stratum pyramidale to the OML2 region of the bottom blade of the DG in a 5–6-month-old adult mice. LFPs were measured using a 64-channel laminar silicon probe (interelectrode distance=20 um; H3, Cambridge NeuroTech, Cambridge, UK). Dorsal hippocampus was identified bilaterally at -2.2 mm AP -1.4 mm ML relative to Bregma. The probe, attached to a digital multiplexing head-stage, was slowly lowered in 1 mm intervals at ∼0.05 mm/s every 10-15 min until the upper channels of the probe (Chs 1-8) were in the CA1 pyramidal cell layer and the bottom probe channels were in or beyond the OML2 region of the DG bottom blade. Probe depth was initially measured relative to skull surface and brain surface (i.e., when first channels at the tip of the probe were grounded against brain signal and free of 60 Hz noise). Continuously sampled LFP recordings were made via the DigitalLynx SX acquisition system and Cheetah software (v6.4, Neuralynx, Boseman, MT) for a minimum of one hour to ensure that cell firing activity and EEG quality were stable. After stereotaxic depth measurements from skull and brain surface, probe position was secondarily determined relative to the CA1 *stratum pyramidale* (SP). Experiment-to-experiment position variability across animals was limited by determining the location of SP and single unit action potentials, and phase reversal from oscillations in descending anatomical regions of hippocampus. As in prior studies, SP location was defined offline as the channel with the maximum ripple power (Dvorak et al., 2021).

Our *in vivo* electrophysiology study primarily focused on network level measures or mesoscale level mechanisms that related network level oscillations to the firing properties of DG granule cells, as a function of DG dendritic compartments. We chose this focus because the high-density laminar probes approach captures a limited number of DG granule cells, where we expect about 13% of cells will be transfected in the *Pten* KO and *Pten; Raptor* dKO groups. In contrast, the laminar probes capture a highly detailed view of the entorhinal-DG synaptic inputs at specific dendritic compartments and DG granule cell outputs via somatic action potentials.

While this approach sacrifices high-density cell recording on the medial-lateral axis for high density dorsal-ventral recordings, it also provides significant gains by allowing us to test levels of dendritic efficacy between wild-type mice, *Pten* KO mice and *Pten; Raptor* dKO mice that are central to our hypotheses. The approach also provides throughput measures that can serve as a bridge between the results of single cell patch physiology experiments and behavioral experiments, as well as the higher transduction experiments analyzing network and survivability outcomes using POMC-Cre+ *Pten* KO and *Pten; Raptor* dKO (∼70-80% transfection).

Theta oscillations reflect the alternating frequency of current propagation along the somatodendritic axes of hippocampal dendritic membranes (Brankack et al., 1993; Buzsaki et al., 1986). Each alternating cycle of EC synaptic inputs to the apical dendrites, their propagation to the cell layer, and their return from the cell layer to the apical dendrites following action potentials, represents a fine balance of inhibition and excitation over a theta scale timeframe (i.e., 1/7 Hz = 142 ms). Theta oscillations therefore allow for biophysical measurements of the temporal throughput organization, and putative hypersynchrony, between entorhinal inputs and dentate gyrus granule cells, and their reversal from cell layer to synaptic inputs. We hypothesize that mutations in the mTOR pathway, for either *Pten* KO or *Pten; Raptor* dKO mice, will affect the activity of granule cells and how dendrites receive and process inputs from the entorhinal cortex. The efficacy of this pathway will be measured through via the quality of theta currents generation.

The entorhinal cortex projects to the hippocampus via the trisynaptic loop (L2 MEC/LEC → DG, DG → CA3, CA3 → CA1) and direct projections from L3 MEC to the CA1 apical dendritic tufts in SLM (Hafting et al., 2005; Sargolini et al., 2006; Witter and Moser, 2006). The circuit is completed with CA1 projections back to L5 of the EC (Ohara et al., 2023). Wheel running reliably shifted the hippocampal circuit from spontaneous activity to robust 5-12 Hz oscillations. Signals were largest in regions corresponding to medial EC layer III and layer II inputs to CA1 and DG. We used theta oscillations to assay signal coordination and the efficacy of coding mechanisms along the neuronal axis of DG neurons. To address our hypotheses, we make 2 primary local field potential measures as a function of depth along the CA1 and DG axes with respect to theta oscillations, or inter-ictal epileptiform discharges in the case of a subset of PTEN KO mice: 1) Voltage rasters; and 2) Current Source Density (CSD). Lastly, we analyze the theta phase preference of excitatory DG granule cells as a function of theta oscillations at 4 locations along the DG somatodendritic axis.

Inter-ictal Epileptiform Discharges: IEDs were detected using a line-length transform algorithm across the 64-channel probe array that was adapted for high-throughput batch processing. The raw voltage signal *d* was transformed into a line-length metric *L* over a sliding window of 40 ms (*llw*), capturing the signal’s path length to emphasize high-frequency, high-amplitude events typical of IED spikes. Events were identified when the line-length signal exceeded a 99.9th percentile threshold (*prc*) calculated from the data distribution. Detections occurring within 300ms of each other were merged into single events to prevent double-counting of multiphasic waveforms. A minimum duration threshold of 25 ms was applied to exclude transient artifacts. The detector generated two primary matrices for each session, event start/stop indices and a binary matrix indicating which channels participated in each event. Following automated detection, candidate events underwent a manual curation process to separate genuine IEDs from other physiological events or artifacts. A custom visualization tool extracted a ± 50 ms window around the midpoint of each detected event. Waveforms across all channels were stacked and plotted to visualize the event’s spatiotemporal profile. Trained observers manually reviewed these visualizations using a Python-based curation interface. Events were classified based on their morphological signature where interictal spikes characterized by a sharp, high-amplitude transient event followed by a slow wave that were visible across multiple adjacent channels were considered for further analysis (Figure S9).

All downstream analyses were orchestrated by a custom MATLAB pipeline (Figure S10) that managed the data iteration through subject directories, ensuring that the same algorithmic standards were applied to every recording session without manual intervention. For each session, the pipeline automatically ingested the pre-processed voltage matrix and the corresponding event timestamp file. Legacy event files were automatically converted to a standardized tabular format prior to analysis. The pipeline sequentially triggered seven independent analytical modules (Theta/Waveform Raster, Event Triggered Waveform Stacks, Voltage Raster, CSD Raster, CSD Center Slices Initiation Phase, CSD Time-Averaged Slices Propagation Phase, and Time-Frequency Scalograms). Each module operated within an error-handling framework to prevent localized artifacts from halting the batch processing of an entire dataset. Following analysis, statistical outputs (e.g., peak amplitudes, half-widths) from all modules were aggregated into a master CSV file and summary visualizations were compiled into high-resolution graphics for rapid quality control and figure generation.

Voltage Raster: For IED analysis, we analyzed event-triggered voltage traces across all recording channels using an Event Stacks module. Precise temporal alignment was critical for averaging multi-trial data. Although events were initially detected via the line-length metric, this method can introduce minor temporal jitter relative to the waveform peak. To correct for jitter, the pipeline performed a local peak search within a narrow window (±5 ms) around the initial detection timestamp. The “common anchor” for each event was defined as the sample index of the maximum absolute voltage deflection on a reference channel. This was typically the channel with the highest signal-to-noise ratio or the deepest channel, depending on the specific run configuration. Voltage traces were segmented into 100 ms windows (±50 ms) centered on this refined anchor point. Events where the peak search failed or the window extended beyond the recording duration were automatically excluded from the average.

Individual event traces were averaged point-by-point to generate a Mean Voltage Matrix (Channels × Time) for each group. This averaging process suppresses incoherent noise and highlights the robust laminar features of the epileptiform discharge. The resulting matrix was rendered as a heatmap over a ± 20 ms window. Voltage amplitude was mapped by color (Jet colormap), with warm colors indicating positive deflections and cool colors indicating negative deflections. To ensure visual comparability between groups, the color limits (CLim) were fixed globally based on the 99.5th percentile of the combined voltage distribution. The raw voltage values of the mean matrix were exported to CSV files to facilitate secondary statistical testing or external plotting.

Current Source Density: The spatial and temporal distribution of local synaptic currents was assessed by current-source density analysis (CSD) (Brankack et al., 1993; Kloc et al., 2023). LFPs were filtered for theta oscillations (4-12 Hz) and CSD was estimated as the second spatial derivative along the 64-channel laminar silicon probe. For each channel k, the CSD signal Ck was derived using the standard three- point finite difference formula: Ck = −(Vk−1 − 2Vk + Vk+1), where V is the voltage potential. Because the spatial derivative requires adjacent channels, the most superficial (Channel 1) and deepest (Channel 64) contacts could not be computed and were assigned NaN values, effectively excluding them from the analysis. A random 6 sec theta epoch during wheel running was selected for CSD analysis in each animal from each group. The current sinks and sources were then isolated across the theta epoch and then averaged. GEE analyses tested for a Group x Depth interaction for mean current source, and tested for specific location differences (i.e., GCL2).

For IED analysis, we evaluated spatial consistency of the current sinks and sources at the precise moment of IED onset as the “Initiation Phase” of the CSD or Voltage raster. Using the CSD matrices, we extracted a 1 ms temporal slice (averaging from 1 ms to 0 ms relative to the anchor) for every IED event. This short averaging window was applied to minimize the impact of single-sample noise while preserving the instantaneous laminar structure of the discharge peak. The spatial profiles were visualized as CSD amplitude (x-axis) shown as a function of channel depth (y-axis). Individual trial profiles were overlaid in semi-transparent gray to visualize variability, while the ensemble mean profile was overlaid in bold black to identify the robust location of the primary sink (negative deflection) and return sources (positive deflection).

To characterize the spatiotemporal evolution of the discharge after the initial peak, we analyzed the “Propagation Phase” of the CSD and voltage raster. Unlike the instantaneous “Initiation Phase” (which captures the peak at t = 0), this analysis integrated the signal over the 2 ms to 15 ms window relative to the anchor point. This window was selected to capture the spread of activity and the subsequent return currents (repolarization or synaptic inhibition) following the primary discharge. For each event, the CSD matrix was averaged across this specific time window to produce a single spatial vector representing the sustained laminar profile of the initial IED event’s aftermath.

The Grand Average was computed as the arithmetic mean of the subject-level matrices:

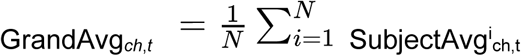

where N is the number of animals in the group. This “Average of Averages” approach ensures that each animal contributes equally to the final result, preventing subjects with higher event counts from skewing the group morphology.

Anatomical boundaries were manually identified for each recording session using a custom Python graphical interface. The 64-channel linear probe data were visualized as Theta-band (4-12 Hz) overlays next to Current Source Density CSD heatmaps. Using these electrophysiological landmarks, specifically the theta phase reversal at the hippocampal fissure and the location of the primary evoked sink, each channel was assigned to a specific hippocampal subfield. The probe trajectory was segmented into 8 distinct internal physiological regions representing the laminar structure of the hippocampus: CA1 Stratum Lacunosum-Moleculare (SLM), followed by the dorsal blade of the Dentate Gyrus Outer Molecular Layer 1, Middle Molecular Layer 1, Granule Cell Layer 1, the Hilus, and the ventral blade Granule Cell Layer 2, Middle Molecular Layer 2, Outer Molecular Layer 2. This approach corrects for anatomical jitter, ensuring that the averaged signals reflect activity from homologous laminar structures across all subjects.

Theta Phase Preference Analysis: Local field potentials (LFPs) were derived from Neuralynx continuous sampling channel (CSC) recordings. For all analyses, raw CSC signals were loaded directly using the Neuralynx MATLAB API (Nlx2MatCSC) by extracting sample values only (FieldSelectionFlags = [0 0 0 0 1]). Signal polarity was inverted to maintain consistency with prior analyses and hippocampal laminar conventions. Raw data were assumed to be sampled at 30 kHz. The signal processing pipeline for theta phase preference beyond this point was modeled after prior work from the Buzsaki lab (Valero et al., 2022).

To obtain LFP signals suitable for oscillatory analysis, raw CSC signals were downsampled to 1,250 Hz using polyphase finite impulse response (FIR) resampling (resample, MATLAB). This procedure includes an implicit anti-aliasing low-pass filter and avoids spectral folding artifacts. Time vectors were reconstructed from the resampled data assuming uniform sampling. Theta epochs were detected once per recording session using a designated reference channel specified a priori for each session. This reference channel was used exclusively for defining global theta periods applied to all units and channels within that session.

Theta detection was performed using a complex Morlet wavelet transform (cwild-type, MATLAB) applied to the downsampled LFP. A wavelet-based formulation was chosen to provide frequency-adaptive estimates of theta phase and power without requiring explicit band-pass filtering or Hilbert transformation. This approach is robust to slow fluctuations in theta frequency and ensures methodological consistency between theta detection and phase estimation. Wavelet coefficients were computed across frequencies spanning 4–14 Hz. At each time point, theta-band power was estimated as the root-mean-square (RMS) amplitude across all wavelet coefficients within this frequency band.

A theta power threshold was defined as the session-wide mean theta power plus 0.8 standard deviations. Time points exceeding this threshold were labeled as theta-active. Consecutive theta-active samples were merged into contiguous theta epochs. To exclude brief or spurious detections, only theta epochs with a minimum duration of four theta cycles at the upper band limit (14 Hz; minimum duration ≈ 0.286 s) were retained. The resulting theta epochs were stored as start–stop time intervals and used as a global gating signal for all subsequent spike–phase analyses within that session. Instantaneous theta phase was estimated independently for each LFP channel using the same wavelet-based approach. For each channel, the downsampled LFP signal was transformed using a complex Morlet wavelet across the 4–14 Hz frequency range. At each time point, the theta frequency with the maximal instantaneous amplitude was identified, and the phase of that wavelet component was extracted. This approach allows the phase estimate to adapt to slow fluctuations in theta frequency rather than assuming a fixed center frequency. Theta phase values were shifted by π radians such that the trough of the theta cycle corresponded to 0° and the peak to 180°, consistent with standard hippocampal phase conventions. The resulting phase time series was stored at the LFP sampling rate for subsequent interpolation.

Unit activity was filtered from LFP recordings and clustered using the automated spike-sorting software Kilosort4 (Pachitariu et al., 2024). All units were examined using Phy 2.0. Units were only considered for analysis if their clusters were considered ‘good’ by Phy: 1) Represent a well-isolated single unit; 2) The auto-correlogram (ACG) shows a significant central dip to approximately zero for a period of 1 ms or more, indicating a clear refractory period where the neuron cannot fire a second action potential; and 3) The spike waveforms are consistent, stable over time, and clearly distinct from other clusters in feature space and amplitude views. Beyond Phy, extracellular DG action potentials were only included in subsequent analysis if they exhibited: 1) An average unit amplitude >50 µV; and 2) A wide waveform shape and peak-trough width consistent with excitatory neurons (> 300-350 µs). Narrow waveform spike clusters (peak-trough time of <350 ms) identified in the cell layer were designated as “likely inhibitory” and excluded from analysis (Fox and Ranck, 1981; Robbins et al., 2013) (Figure S11).

Spike timestamps for individual clusters were loaded from precomputed timestamp files and converted from sample indices to seconds. For each spike, instantaneous theta phase was obtained by linear interpolation of the unwrapped wavelet phase time series at the spike time, followed by re-wrapping to the [−π, π] range. Spike phases were then gated by the session-wide reference theta epochs. Only spikes occurring within reference-defined theta intervals were included in phase-locking analyses; spikes outside theta periods were excluded.

Statistics for behavior and in vivo electrophysiology: Experimenters were blind to treatment group throughout all experiments. The blind was broken after behavioral acquisition and most electrophysiological analyses were complete. All data described are presented as mean ± standard error (SE). For all statistical tests, p<0.05 was considered significant. For spatial accuracy behavior, the outcome measure is proportion time in goal zone. Generalized linear models (GLM; IBM SPSS v26.0, Armonk NY) or generalized estimating equations (GEE) (Ziegler et al., 1998) were used to generate Wald values to determine significant differences within and between groups. For behavior, GLM and GEE were used to test for interactions of effect with training phase (Phases 7-10). For in vivo electrophysiology, we tested for group interactions with depth location or anatomical region (i.e., GCL2) for theta oscillations or IEDs. Models were adjusted for repeated measures according to the distribution of each analyzed variable (i.e., log link models were used for nonparametrically distributed data). Goodness of fit was determined using the corrected quasi likelihood under independence model criterion.

For phase preference analysis, circular statistics were evaluated using Rayleigh’s test for non-uniformity of nonlinear data (Fisher, 1995) within the Circular Statistics Toolbox in MATLAB (Berens, 2009a). Theta was sampled from contacts at each third of the somatodendritic axis for cells recorded on the suprapyramidal blade (GC1) or infrapyramidal blade (GC2), as defined by depth measurements and CSD analysis. The results were then pooled for statistical analysis.

Finally, we used Receiver Operating Characteristics curves (ROC) to analyze the diagnostic ability of a binary classifier (SPSS 27.0; Armonk, NY). The True Positive Rate (TPR) or sensitivity is plotted against the False Positive Rate (FPR), calculated as 1-specificity, at various threshold settings. We used high percentage bilateral PTEN KO rate as a binary state measure of the number of training trials to criterion, or the PTEN KO positive IED mice as a binary state measure of the mean theta CSD per anatomical region. We found that high bilateral PTEN KO was a predictor of behavioral outcomes in learning to associate a goal object with food reward, and that mean GCL2 and MML2 theta CSD measures were a predictor of IEDs.

The Area Under Curve (AUC) value is used to determine the ability of the test measure to predict cognitive outcome. An AUC value at 0.50 is statistically no better than chance, 0.51-0.69 is considered a poor predictor, 0.70-0.89 is considered a good predictor, and 0.90-0.99 is considered an excellent predictor (Carter et al., 2016).

### Video/EEG Monitoring

EEG implant surgery was performed in mice at 6–9 weeks of age. Animals were anesthetized with 4% isoflurane and positioned in a stereotaxic apparatus using ear bars. Anesthesia was maintained at 1.5–2.5% isoflurane throughout the procedure. The skull surface was exposed, and burr holes were drilled at four locations using a stereotaxic-mounted drill. Stainless steel 3/32″ screws were inserted epidurally to serve as EEG recording electrodes.

Screws were threaded through a head mount and secured using VetBond followed by conductive epoxy. Recording electrodes were placed bilaterally at approximately the following coordinates relative to bregma: AP +1.0 mm, ML ±1.0 mm; AP −1.0 mm, ML ±1.5 mm; and AP −2.5 mm, ML ±1.5 mm. EMG leads were inserted into the nuchal muscle. The head mount was further secured to the skull using dental cement. Animals received ketoprofen (5 mg/kg) prior to surgery and were allowed to recover for at least 5 days before EEG recording. Mice were monitored and weighed twice daily for 3 days postoperatively and received additional ketoprofen if signs of pain or distress were observed.

Mice were housed in ventilated cages under controlled environmental conditions (22–23 °C, ∼60% humidity) on a 12 h light/12 h dark cycle (lights on at 7:00 AM, off at 7:00 PM), with ad libitum access to food and water. EEG recordings were obtained using a Pinnacle three-channel EEG/EMG system when mice were 7–12 weeks old. Recordings were continuous for 120 h, sampled at 1 kHz, and hardware-filtered at 400 Hz. Prior to analysis, EEG signals were bandpass filtered between 0.5 and 100 Hz, and EMG signals were high-pass filtered at 10 Hz using Sirenia software (Pinnacle Technology, version 2.2.8).

EEG recordings were reviewed in their entirety by two independent raters. Generalized tonic–clonic seizures (GTCSs) were defined as high-amplitude EEG activity lasting at least 20 s with an evolving waveform pattern. All GTCS events were confirmed by synchronized video to be associated with stage 5 Racine behaviors, including rearing and falling with forelimb clonus. Non-convulsive seizures in *Pten^flx/flx^;Raptor^flx/flx^*x POMC-Cre+ animals were identified as high-amplitude seizure-like EEG events lasting at least 5 s and verified to not be associated with any movement artifacts.

### CME bHPLC phosphoTMT Methods – Orbitrap Eclipse

POMC-Cre- and POMC-Cre+ animals for *Pten^flx/flx^* and *Pten^flx/flx^ Raptor^flx/flx^* were anesthetized with 2% avertin, transcardially perfused with sterilized and ice-cold saline solution (0.9% NaCl). Dentate gyri were dissected (Olympus MVX10) then snap-frozen on chilled ethanol mixed with dry ice. Total protein from each sample was reduced, alkylated, and purified by chloroform/methanol extraction prior to digestion with sequencing grade trypsin and LysC (Promega). The resulting peptides were labeled using a tandem mass tag 11-plex isobaric label reagent set (Thermo), combined into a single multiplex group, then enriched using High-Select TiO2 and Fe-NTA phosphopeptide enrichment kits (Thermo) following the manufacturer’s instructions. Both enriched and un-enriched labeled peptides were separated into 46 fractions on a 100 x 1.0 mm Acquity BEH C18 column (Waters) using an UltiMate 3000 UHPLC system (Thermo) with a 50 min gradient from 99:1 to 60:40 buffer A:B ratio under basic pH conditions, then consolidated into 18 super-fractions for both the enriched and un-enriched sample sets.

Buffer A = 10 mM ammonium hydroxide, 0.5% acetonitrile,

Buffer B = 10 mM ammonium hydroxide, 99.9% acetonitrile

Both buffers adjusted to pH 10 for offline separation

Each super-fraction was then further separated by reverse phase XSelect CSH C18 2.5 um resin (Waters) on an in-line 150 x 0.075 mm column using an UltiMate 3000 RSLCnano system (Thermo). Peptides were eluted using a 75 min gradient from 98:2 to 60:40 buffer A:B ratio. Eluted peptides were ionized by electrospray (2.4 kV) followed by mass spectrometric analysis on an Orbitrap Eclipse Tribrid mass spectrometer (Thermo) using multi-notch MS3 parameters. MS data were acquired using the FTMS analyzer in top-speed profile mode at a resolution of 120,000 over a range of 375 to 1500 m/z. Following CID activation with normalized collision energy of 31.0, MS/MS data were acquired using the ion trap analyzer in centroid mode and normal mass range. Using synchronous precursor selection, up to 10 MS/MS precursors were selected for HCD activation with normalized collision energy of 55.0, followed by acquisition of MS3 reporter ion data using the FTMS analyzer in profile mode at a resolution of 50,000 over a range of 100-500 m/z. Buffer A = 0.1% formic acid, 0.5% acetonitrile Buffer B = 0.1% formic acid, 99.9% acetonitrile

### Data Analysis – (pTMT)

Proteins were identified and reporter ions quantified by searching the UniprotKB *Mus musculus* database (Proteome ID: UP000000589, 3^rd^ version of 2025) using MaxQuant version 2.2.0 (Max Planck Institute) with a parent ion tolerance of 3 ppm, a fragment ion tolerance of 0.5 Da, a reporter ion tolerance of 0.001 Da, trypsin/P enzyme with 2 missed cleavages, variable modifications including oxidation on M, Acetyl on Protein N-term, and fixed modification of Carbamidomethyl on C. Protein identifications were accepted if they could be established with less than 1.0% false discovery. Proteins identified only by modified peptides were removed. Protein probabilities were assigned by the Protein Prophet algorithm (Nesvizhskii et al., 2003). TMT MS3 reporter ion intensity values are analyzed for changes in total protein using the unenriched lysate sample. Phospho(STY) modifications were identified using the samples enriched for phosphorylated peptides. The enriched and un-enriched samples are multiplexed using TMT11-plex batches, one for the enriched and one for the un-enriched samples.

Following data acquisition and database search, the MS3 reporter ion intensities were normalized using ProteiNorm (Graw et al., 2020). The data was normalized using Cyclic Loess (Ritchie et al., 2015) and analyzed using ProteoViz to perform statistical analysis using Linear Models for Microarray Data (limma) with empirical Bayes (eBayes) smoothing to the standard errors. Limma was also used for differential analysis. Proteins with an FDR-adjusted p-value < 0.05 and an absolute fold change > 2 were considered significant.

### Bioinformatic Analyses

Proteins with significant differential phosphorylation in *Pten* KO versus wild-type (|log2 fold change| > 1.1, p < 0.05, nominal) that also showed significant differential phosphorylation in *Pten;Raptor* dKO versus wild-type at the same threshold were assigned to the mTORC1-independent (AKT/mTORC2) program (302 proteins). Proteins that were significant in *Pten* KO versus wild-type but not in *Pten;Raptor* dKO versus wild-type were assigned to the mTORC1-dependent program (599 proteins). Before GO analysis, we confirmed that the majority of regulated phosphoproteins (781/901, 87%) showed no corresponding significant change in total protein abundance (p > 0.05 or |log2 fold change| < 0.5 in the paired total proteome dataset), indicating that the regulated phosphopeptides reflect phosphorylation state changes rather than protein expression changes.

Gene ontology enrichment analysis was performed using g:Profiler (version e114_eg62_p19; Raudvere et al., 2019) with the Mus musculus annotation database (PANTHER version 19). We queried GO Biological Process, GO Molecular Function, GO Cellular Component, KEGG, and Reactome databases simultaneously. Statistical significance was assessed using g:Profiler’s g:SCS multiple testing correction, with a threshold of FDR < 0.05. For the enrichment background, we used the union of all unique proteins detected in this experiment: all proteins quantified by total TMT proteomics (5,500 proteins) and all proteins identified in the phospho-enriched TMT dataset (3,371 proteins), yielding a tissue-matched background of 6,446 unique dentate gyrus proteins. This approach was preferred over the full annotated mouse genome to avoid artifactual enrichment of neuronal and synaptic terms that arise from any brain-derived protein set compared against a genome-wide reference. Enrichment is therefore reported relative to the detectable dentate gyrus proteome from the same animals and tissue preparation. Ontological enrichment of differentially regulated protein lists was conducted using the gprofiler2 package (v0.2.3)(Kolberg et al., 2020). Protein-protein interactions of the 302 proteins differentially regulated in both *Pten* KO and *Pten;Raptor* dKO were queried from the STRING database (Szklarczyk et al., 2023) using the stringApp (v2.1.1) within Cytoscape (version 3.10.2), with a confidence cutoff of 0.4. The top 30 hub genes were identified using the Maximum Clique Centrality (MCC) method within the cytoHubba app (v0.1)(Chin et al., 2014).

Enrichment of differentially regulated protein lists among disease gene lists was conducted using a hypergeometric test from the hypeR package (v2.2.0) (Federico and Monti, 2020). The background was defined as all unique proteins detected and quantified in the phosphoproteomics experiment. The ASD gene list was obtained from (Fu et al., 2022). The epilepsy gene list was obtained from the Genes4Epilepsy database (Oliver et al., 2023). All disease lists were converted to mouse genes using the biomaRt package (v2.60.1) (Durinck et al., 2009) and were filtered to only genes corresponding to proteins detected and quantified in the phosphoproteomics experiment. All p-values were corrected for multiple hypotheses using the FDR method. Additional gene list comparisons and tables of overlapping genes were made using custom python scripts generated with Claude Code Opus 4.7 using direct uppercase string matching between the human gene symbols in the ASD/epilepsy databases and the PTEN phosphoprotein gene list.

## FIGURES

**Figure S1.**
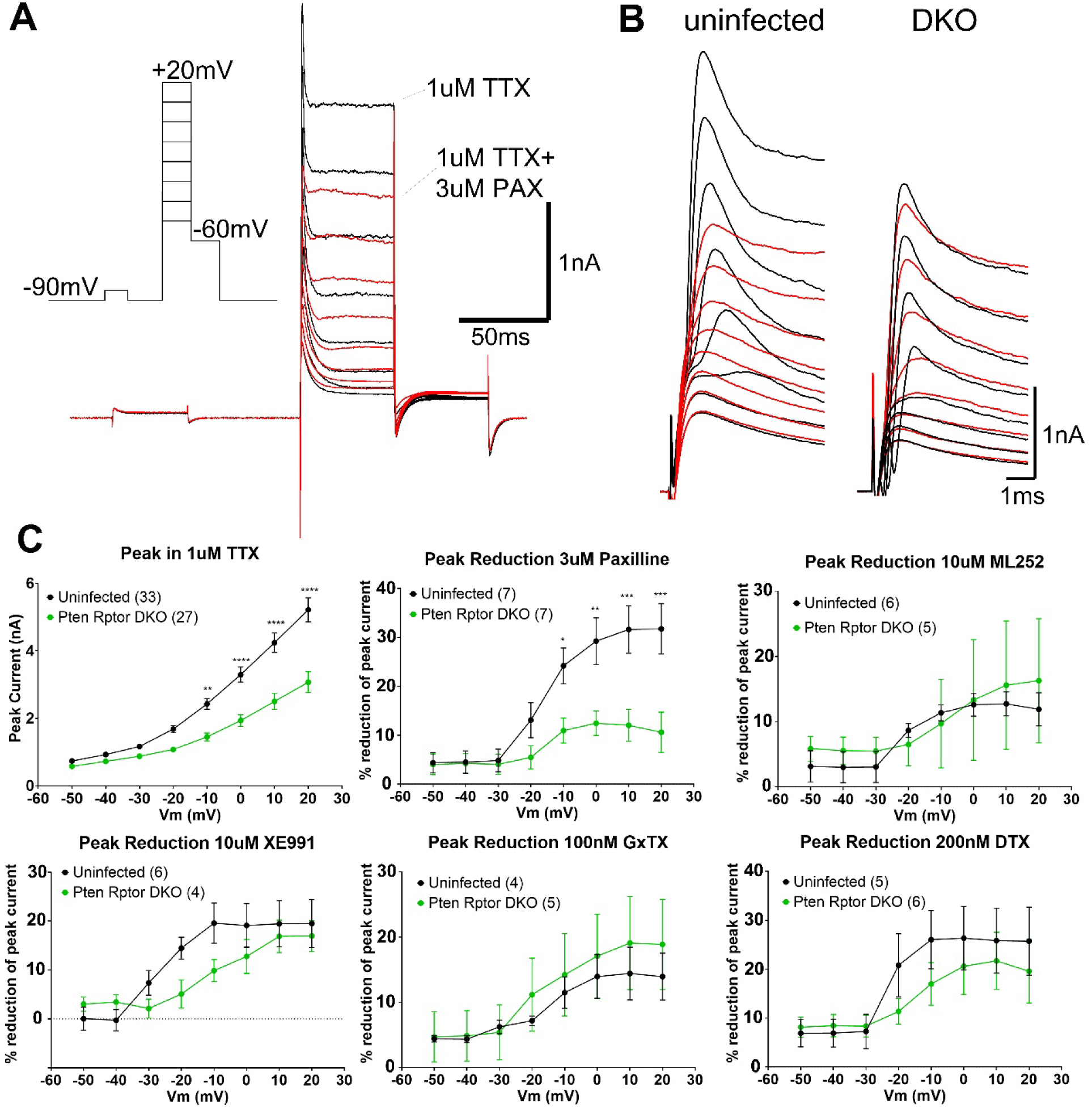
Voltage-Gated K^+^ Currents Are Reduced in *Pten;Raptor* dKO and Are Less Sensitive to K_Ca_1.1(BK) Block. (A) Granule neurons were voltage-clamped at −90 mV and stepped progressively to +20 mV (protocol inset) to evoke outward K^+^ currents (black trace). In uninfected wild-type neurons, 3 µM paxilline (PAX) partially blocked the outward current (red trace). (B) Expanded view of peak K^+^ currents in uninfected control and Cre-expressing *Pten;Raptor* dKO neurons (black traces). Following application of 3 µM paxilline (red traces), peak current was markedly reduced in uninfected cells but largely unchanged in *Pten;Raptor* dKO neurons (compare pre- and post-paxilline traces). (C) Peak K^+^ current amplitude plotted as a function of membrane voltage (Vm) shows reduced current in *Pten;Raptor* dKO neurons (measured in 1 µM TTX). In wild-type neurons, paxilline (3 µM) reduced peak current by ∼30%, compared to ∼5% in *Pten;Raptor* dKO neurons. This genotype-dependent effect was not observed with the K_V_7 blockers ML252 (10 µM) or XE991 (10 µM), the K_V_2 blocker guangxitoxin (GxTx, 100 nM), or the K_V_1 blocker α-dendrotoxin (DTX, 100 nM). Cell numbers are indicated in parentheses; data are presented as mean ± SEM. (*p<0.05, **p<0.01, ***p<0.001 versus wild-type; mixed-effect model with Sidak’s post-hoc).

**Figure S2.**
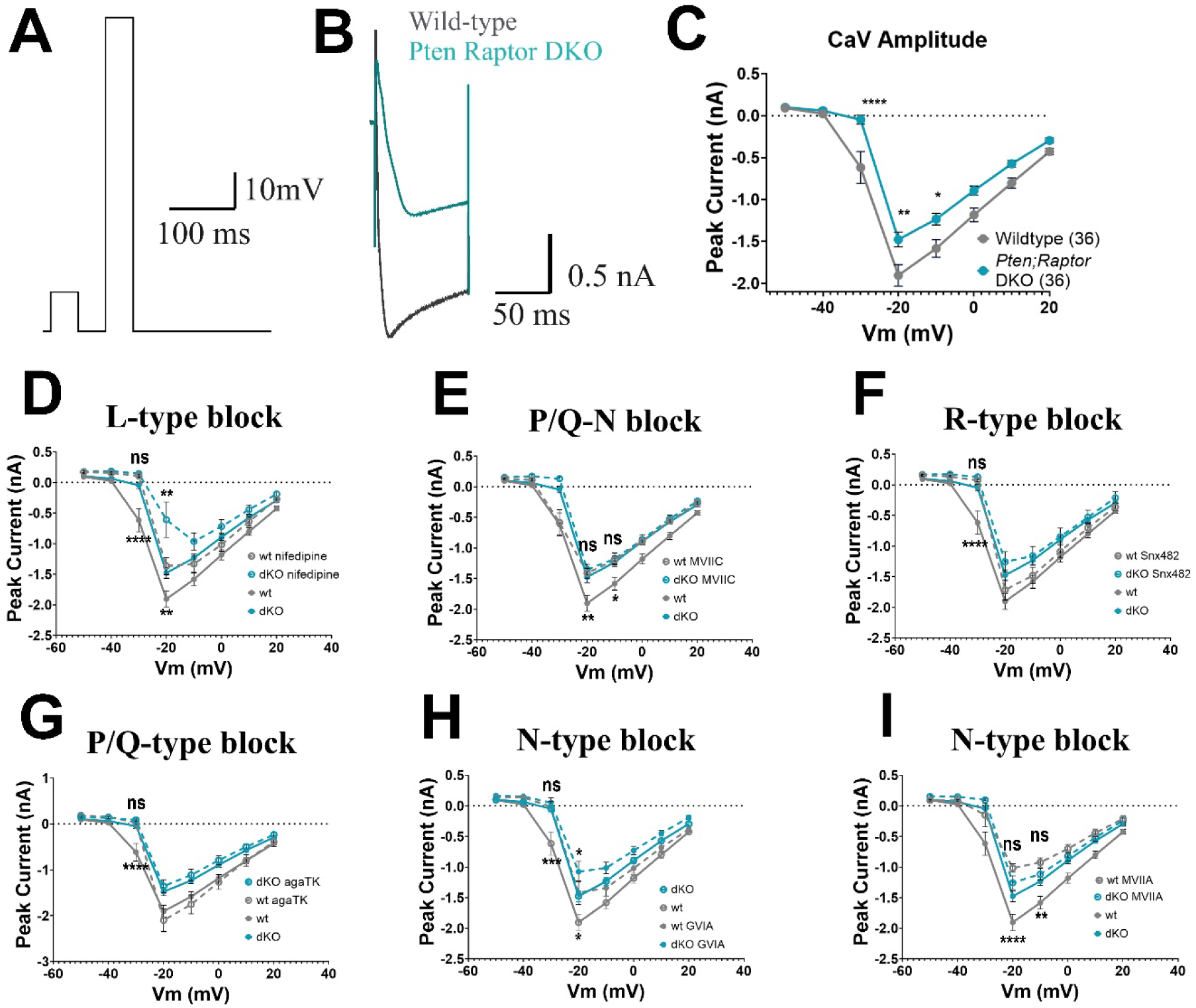
Voltage-Gated Ca Currents are Reduced in *Pten;Raptor* dKO. (A) Voltage-clamp protocol for CaV currents. Neurons were held at −90 mV, followed by a 10 mV test pulse and a series of depolarizing steps from −50 to +20 mV. A representative trace from a −20 mV step is shown. (B) Representative Ca^2+^ current traces from average wild-type (gray) and *Pten;Raptor* dKO (cyan) neurons in response to the −20 mV step. (C) Current–voltage (I–V) relationship plotting peak Ca^2+^ current amplitude (nA) versus membrane voltage (Vm, mV) for wild-type (gray) and *Pten;Raptor* dKO (cyan) neurons. *Pten;Raptor* dKO neurons exhibit reduced voltage-gated Ca^2+^ currents. n values (neurons) are indicated in parentheses (*p<0.05, **p<0.01, ***p<0.001. ****p<0.0001; mixed-effects model with Šidák multiple-comparisons test). (D-I)Comparison of Ca^2+^ currents in wild-type (gray) and *Pten;Raptor* dKO (cyan) neurons recorded in the absence (solid lines) or presence (dashed lines) of specific Ca^2+^ channel blockers: L-type blocker nifedipine (10 µM; D), P/Q- and N-type blocker ω-conotoxin MVIIC (1 µM; E), R-type blocker SNX-482 (0.5 µM; F), P/Q-type blocker ω-agatoxin TK (0.2 µM; G), N-type blocker ω-conotoxin GVIA (1 µM; H), and N-type blocker ω-conotoxin MVIIA (50 nM; I). Statistical comparisons below each graph indicate wild-type versus wild-type + blocker; comparisons above indicate *Pten;Raptor* dKO versus *Pten;Raptor* dKO + blocker. (ns p>0.05, *p<0.05, **p<0.01, ***p<0.001. ****p<0.0001 using mixed-effect model with Tukey’s multiple comparisons for every condition at ever voltage). If no difference is indicated then there was not a significant effect of that drug on wild-type or *Pten; Raptor* dKO neurons at the indicated voltage.

**Figure S3.**
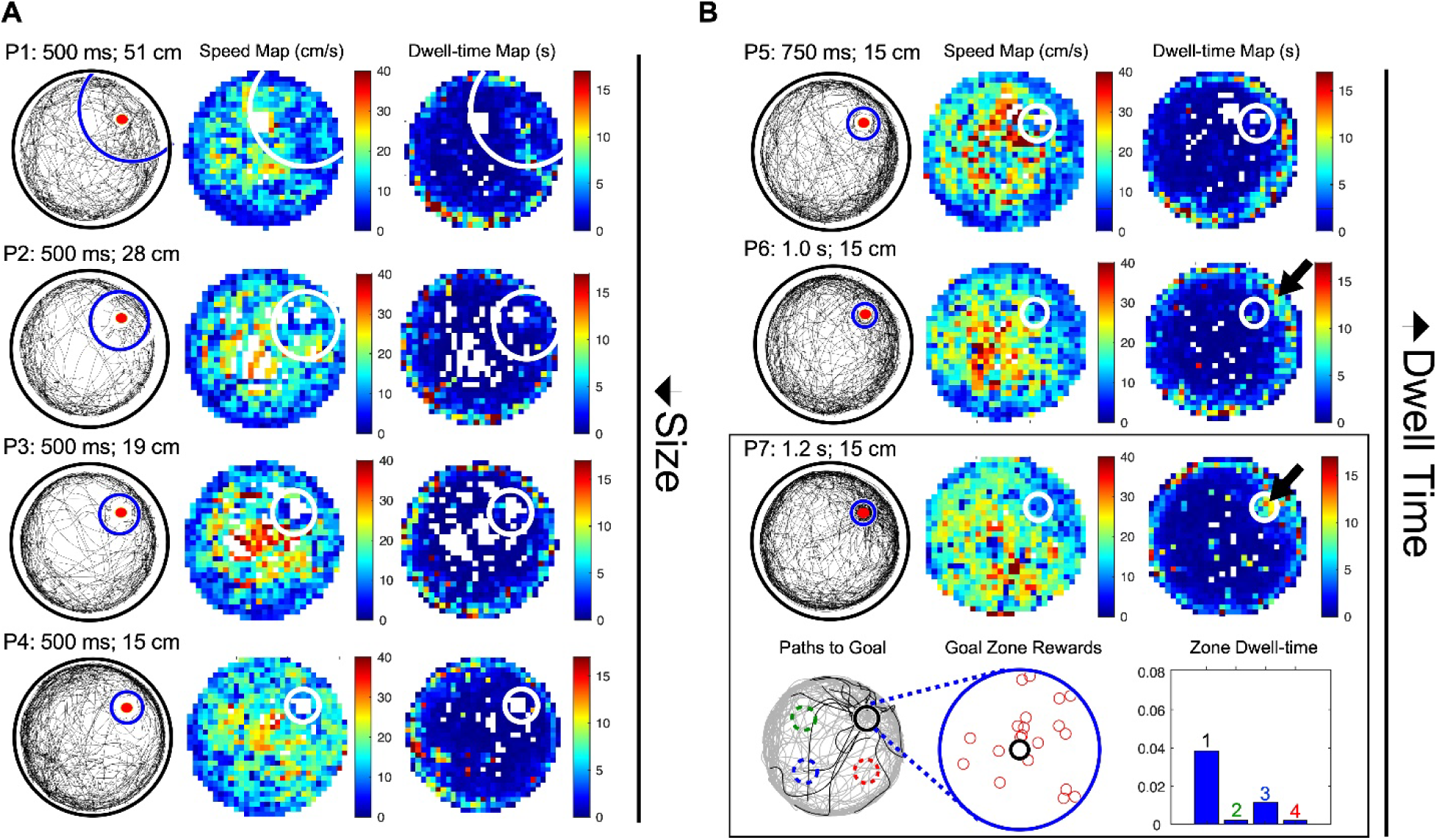
Object Shaping Paradigm and Performance from a *Pten* KO Mouse. (A) Left, overall path (black lines), goal zone (blue semicircle/circle), and goal object (red circle); middle, speed map (cm/s); right, dwell time map. Goal zone for a visible object (V1) diameter progressively decreases from 51 cm to 15 cm during phases P1–P4. (B) Top, required dwell time within V1 increases from 500 ms to 750 ms and 1.2 s during phases P5–P7. Bottom left, rewarded goal entrances (black lines and solid black circle) shown relative to overall path (gray lines) and zones equidistant from the arena center in each quadrant (dashed circles in green, blue, and red). Bottom middle, magnified view of the goal zone showing reward locations. Bottom right, quantification of the proportion of session time spent in zones matching the goal zone diameter (zone 1) and in equidistant control zones in the other three quadrants (zones 2–4). Some *Pten* KO mice required additional sessions to reach criterion due to off-target pauses during early training (black arrow in P6).

**Figure S4.**
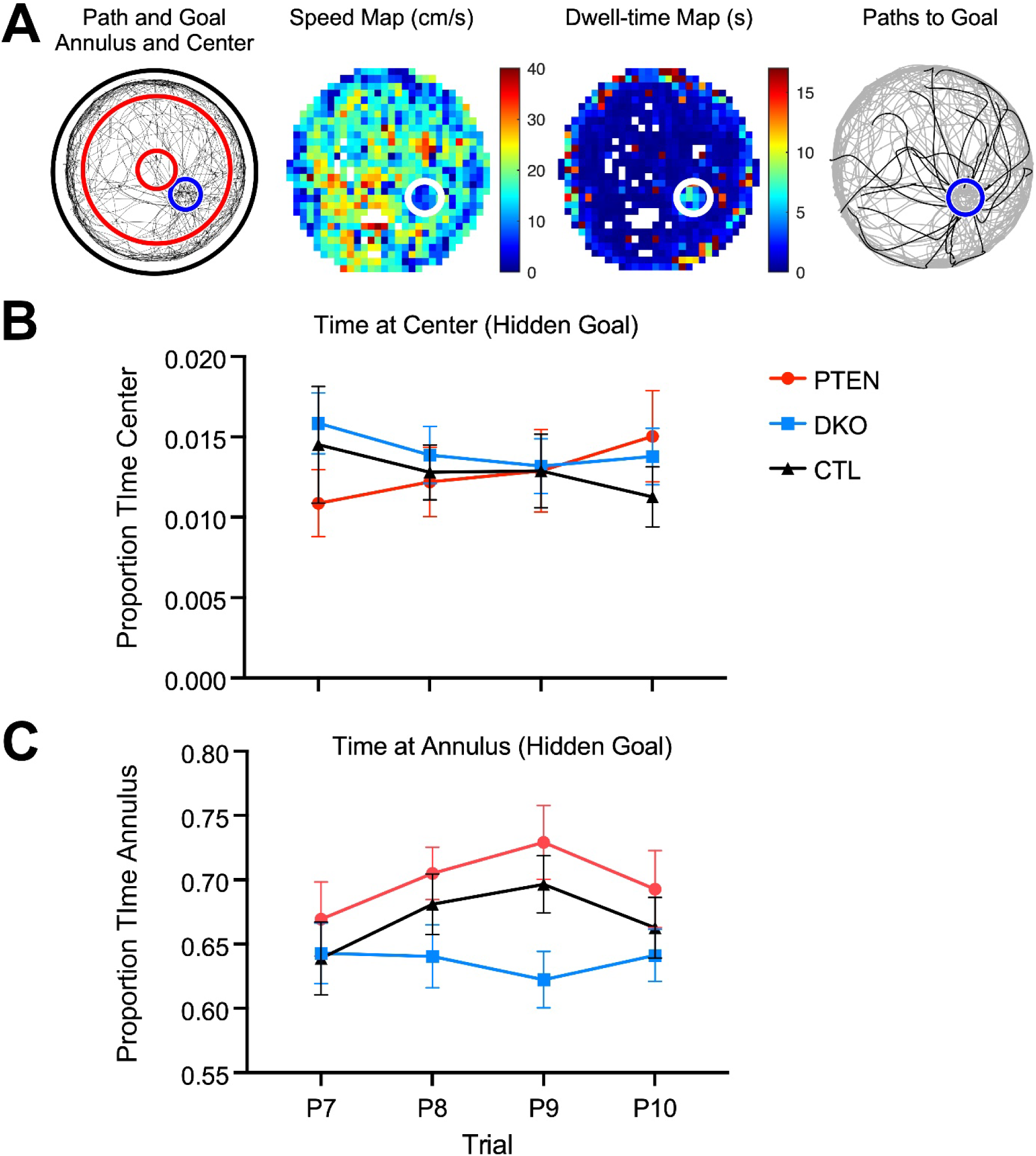
Spatial cognitive differences in *Pten* KO mice are not due to thigmotaxis: (A-C) Measurement of the proportion time (A) spent in the cylinder center (B) or annulus (C) shows no significant group x phase interaction effects. Spatial accuracy deficits shown by *Pten* KO during goal rotation sessions are therefore not due to differences in thigmotaxic behavior or increased anxiety levels.

**Figure S5.**
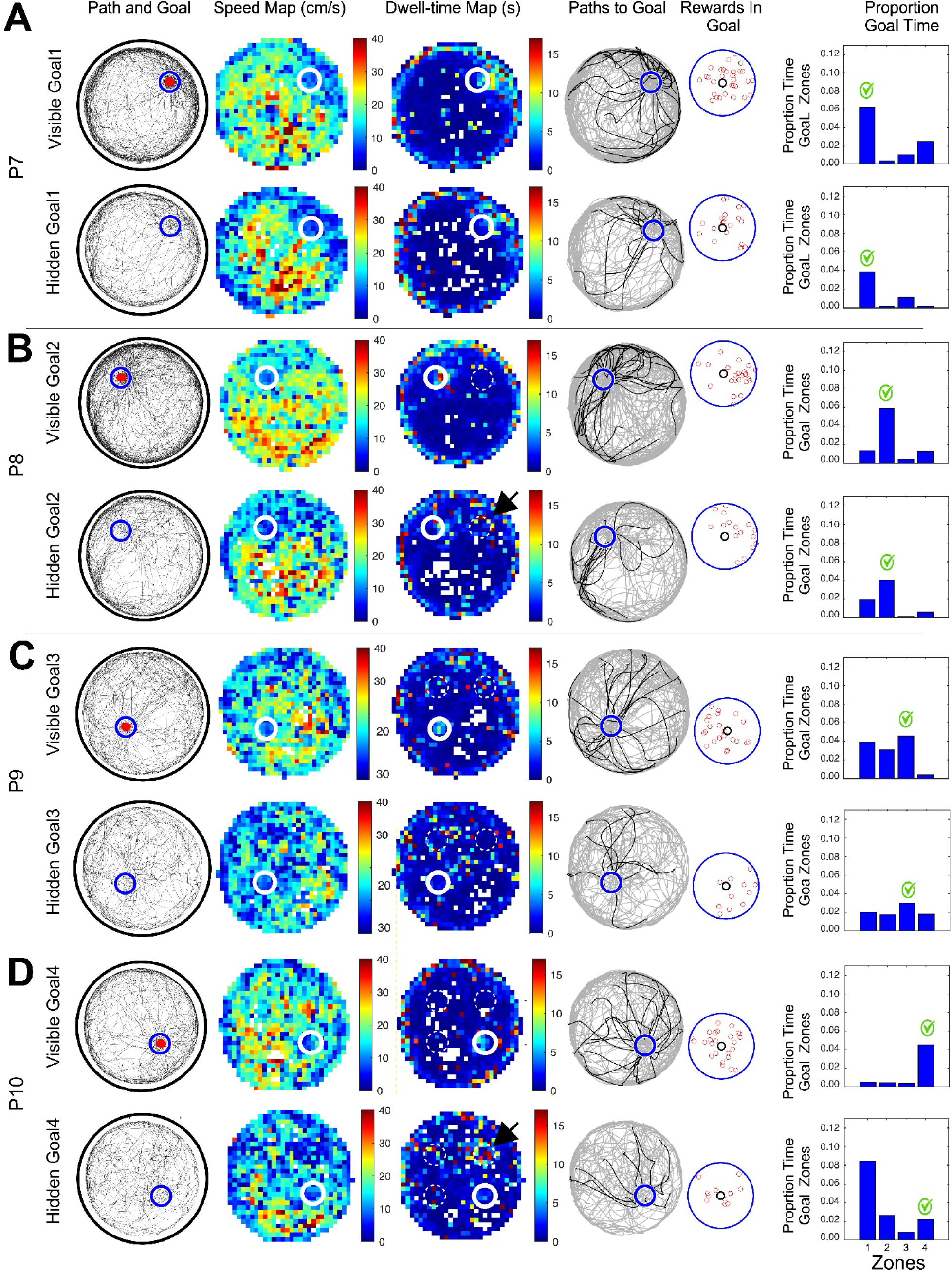
Example performance of a *Pten* KO mouse in visible (V1-4) and hidden goal (H1-4) probe sessions in phases 7-10. As in Figure S2, Performance is shown in 6 plots illustrating the overall path and goal location, speed map, dwell-time map, paths to goal, location of rewards within the goal zone, and quantification of the proportion time spent in equal sized zones the same distance from the arena center in each quadrant. Goal-pauses during visible goal probes were generally restricted to the correct goal zone (but see phase 9). Yet in hidden goal sessions, the *Pten* KO mouse tended to keep pausing in zone 1, particularly during Phase 10 when H4 was the correct goal zone.

**Figure S6.**
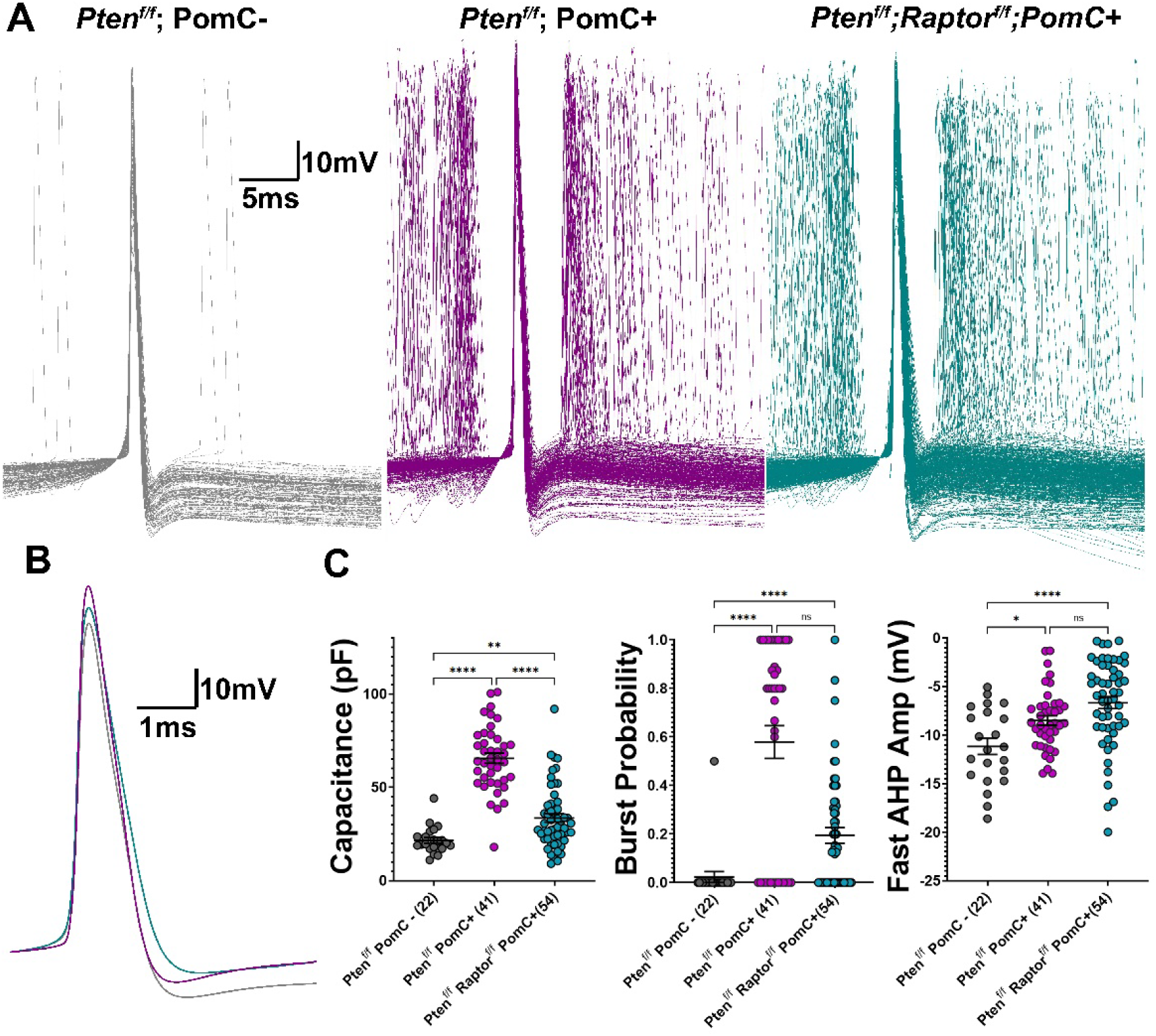
Whole-Cell Electrophysiology in POMC-Cre *Pten* and *Pten;Raptor* dKO Confirms Bursting Phenotypes Found in Retroviral KO. (A) Overlay of action potentials recorded at rheobase from *Pten^flx/flx^;*POMC-Cre− (gray), *Pten^flx/flx^*;POMC-Cre+ (magenta), and *Pten^flx/flx^;Raptor^flx/flx^*;POMC-Cre+ (cyan) neurons, illustrating prominent burst firing in *Pten* KO and *Pten;Raptor* dKO neurons. (B) Average action potential waveforms from the same genotypes showing reduced fast afterhyperpolarization (fAHP) amplitude in *Pten* KO and *Pten;Raptor* dKO neurons. (C) Quantification of membrane capacitance, burst probability, and AHP amplitude across genotypes. (n, is indicated in parentheses; *p<0.05, **p<0.01, ***p<0.001, ****p<0.0001 for capacitance and fAHP a one-way ANOVA with Tukey’s post-hoc and for burst probability a fisher’s exact test with the contingency of burst vs no burst

**Figure S7.**
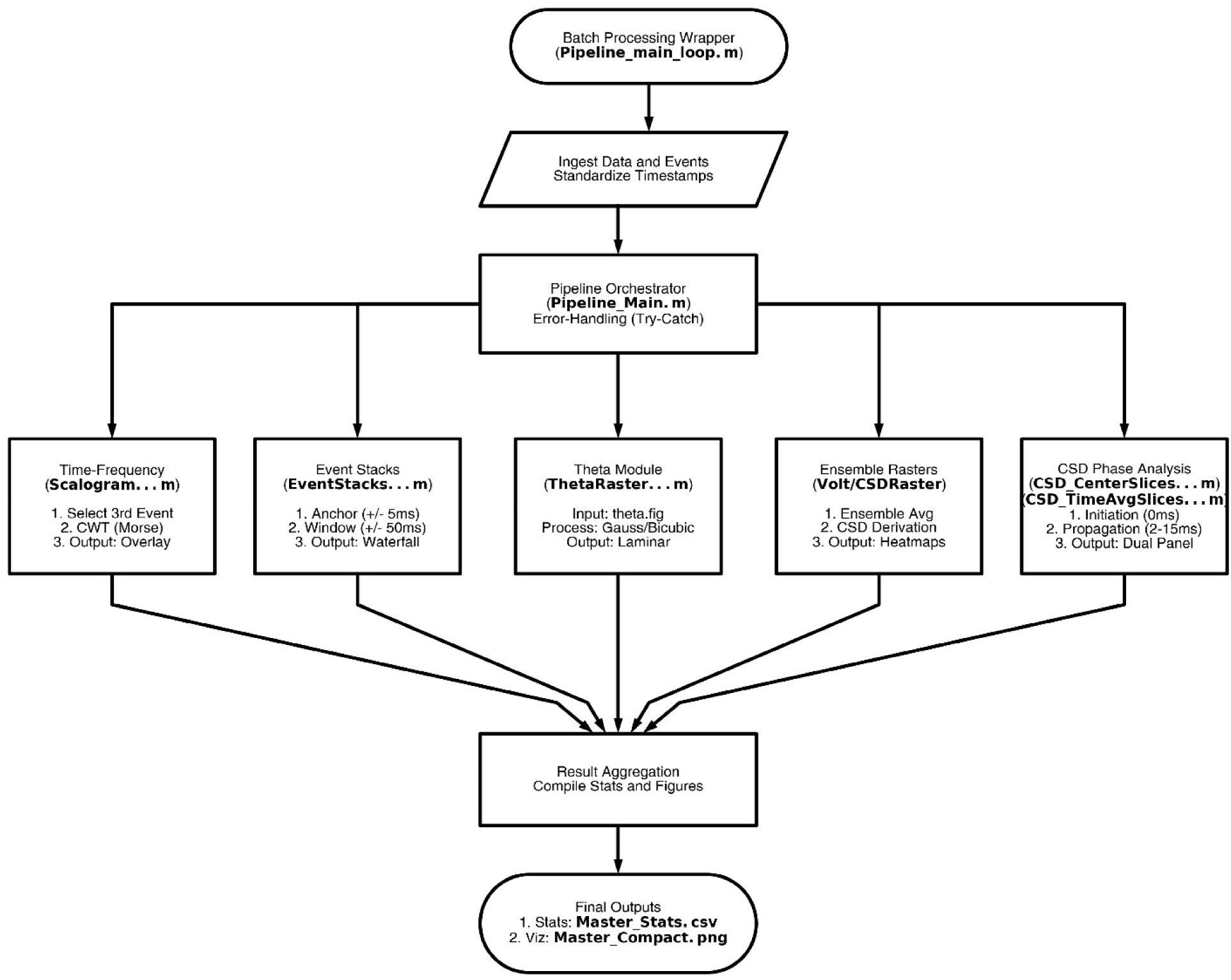
LFP data analysis pipeline. To ensure consistent parameter application and reproducibility across large-scale datasets, all downstream analysis was orchestrated by a custom MATLAB pipeline. A wrapper script managed the iteration through subject directories, ensuring that the same algorithmic standards were applied to every recording session without manual intervention. For each session, the pipeline automatically ingested the pre-processed voltage matrix and the corresponding event timestamp file. Legacy event files were automatically converted to a standardized tabular format prior to analysis. The pipeline sequentially triggered seven independent analytical modules (Theta/Waveform Raster, Event Triggered Waveform Stacks, Voltage Raster, CSD Raster, CSD Center Slices Initiation Phase, CSD Time-Averaged Slices Propagation Phase, and Time-Frequency Scalograms). Each module operated within a robust error-handling framework to prevent localized artifacts from halting the batch processing of an entire dataset Following analysis, statistical outputs (e.g., peak amplitudes, half-widths) from all modules were aggregated into a master CSV file and summary visualizations were compiled into high-resolution figure panels.

**Figure S8.**
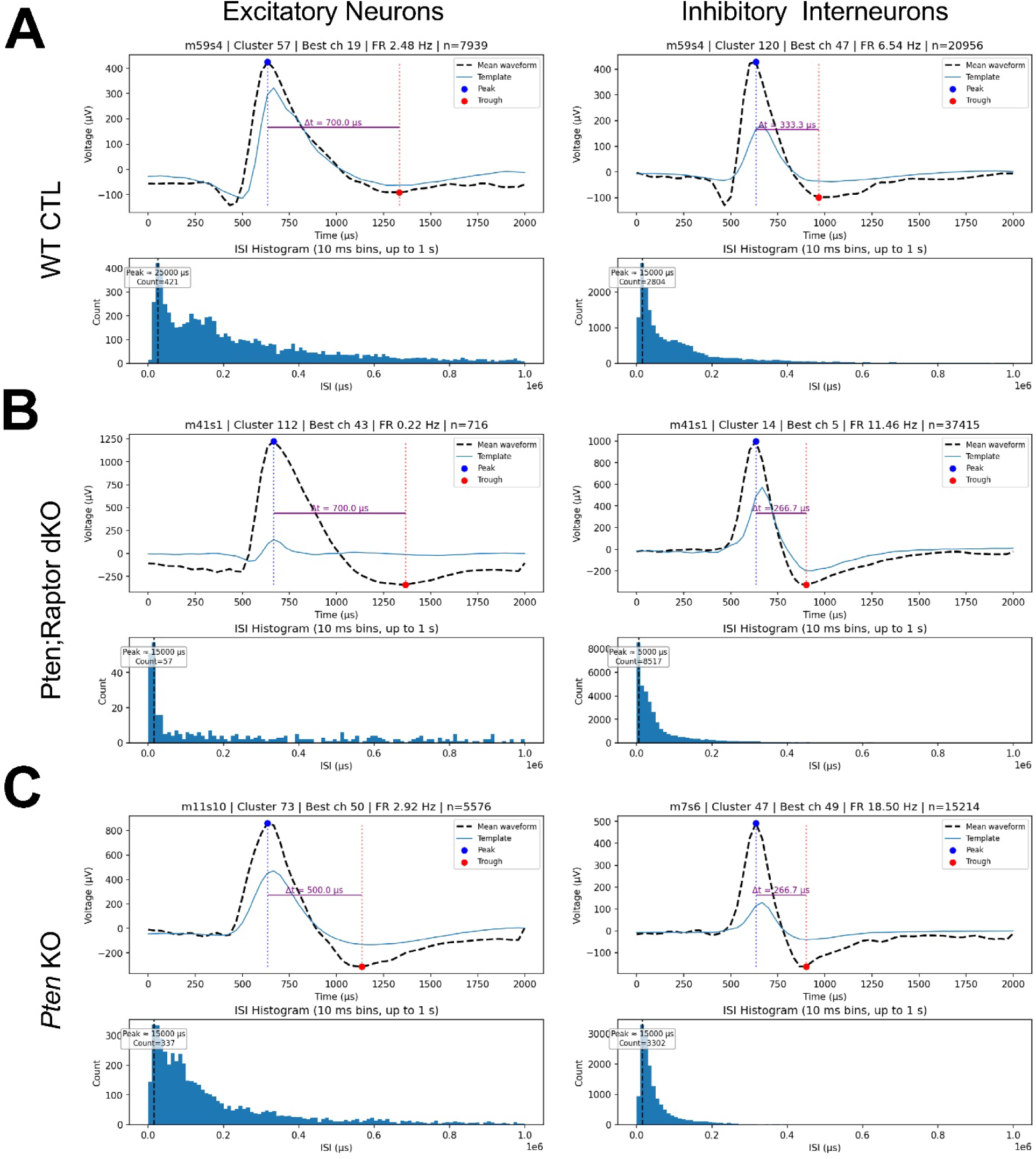
Example Kilosort clustering. (A-C). Classification of excitatory (left) and inhibitory (right) neurons based on extracellular waveform properties. Kilosort was used for automated spike sorting and clustering, followed by analysis of waveform features including spike duration and inter-spike interval (ISI) for wild-type (A), *Pten;Raptor* dKO (B), and *Pten* KO (C) mice.

